# An original potentiating mechanism revealed by the cryoEM structures of the human α7 nicotinic receptor in complex with nanobodies

**DOI:** 10.1101/2023.01.03.522595

**Authors:** Marie S. Prevost, Nathalie Barilone, Gabrielle Dejean De La Batie, Stéphanie Pons, Gabriel Ayme, Patrick England, Marc Gielen, François Bontems, Gérard Pehau-Arnaudet, Uwe Maskos, Pierre Lafaye, Pierre-Jean Corringer

## Abstract

The human α7 nicotinic receptor is a pentameric channel mediating cellular and neuronal communication. It has attracted considerable interest to design ligands for the treatment of neurological and psychiatric disorders. To develop a novel class of α7 ligands, we recently generated two nanobodies named E3 and C4 acting as positive and silent allosteric modulators respectively. Here, we solved the cryo-EM structures of the nanobody-receptor complexes. E3 and C4 bind to a common epitope involving two subunits at the apex of the receptor. They form by themselves a symmetric pentameric assembly that extends the extracellular domain. Unlike C4, the binding of E3 drives an active or desensitized conformation in the absence of orthosteric agonist, and mutational analysis shows a key contribution of a N-linked sugar moiety in mediating E3 potentiation. The nanobody E3, by remotely controlling the global allosteric conformation of the receptor, implements an original mechanism of regulation which opens new avenues for drug design.

## Introduction

Nicotinic acetylcholine receptors (nAChRs) mediate cellular communication in neuronal and non-neuronal cells. In humans, sixteen genes encode for nAChRs subunits that can assemble into homo- or hetero-pentamers, generating a wide diversity of combinations each displaying an unique expression pattern and physiological function^1^. Among nAChRs, the homopentameric α7 has attracted considerable interest with the aim of developing enhancers for the treatment of neurological and psychiatric disorders associated with cognitive decline such as Alzheimer’s disease, schizophrenia, autism or bipolar troubles^2^. The α7-nAChR is also an essential component of the cholinergic anti-inflammatory pathway^3^.

nAChRs are pentameric ligand-gated ion channels (pLGIC). Acetylcholine (ACh) binding to the orthosteric site within their extracellular domain (ECD) promotes the fast transition from a resting closed-channel state to an active open-channel state selective for cations. This is followed by a slower transition to a desensitized closed-channel state. α7-nAChR specific agonists and partial agonists have been developed, some of which display pro-cognitive effects in pre-clinical trials^4^. Positive allosteric modulators (PAMs) binding in the transmembrane domain (TMD) also showed beneficial effects, as exemplified by the Type II PAM PNU-120596 (PNU) which potentiates the ACh-elicited currents and strongly impairs desensitization^5^.

Several human α7-nAChR high-resolution structures have recently been solved by cryo-electron microscopy (cryoEM), in detergent micelles or in protein/lipid nanodiscs^6,7^. α7-nAChR harbours a β-folded ECD carrying the orthosteric site at the interface between subunits, coupled to a TMD with 4 α-helices per subunit (termed M1 to M4) that form the ion channel, and an intracellular domain (ICD), which contains two helices (MA and MX) flanking a highly variable region which is truncated in the constructs used for solving cryo-EM structures. Closed-channel structures in Apo and antagonist (bungarotoxin (Bgt))-bound conditions have been annotated as resting conformations, while those in complex with agonists (epibatidine or EVP-6124 abbreviated as EVP) have been annotated as desensitized conformations. Structures in complex with both agonist and PNU show an apparently open (Epi) or semi-open (EVP) pore, consistent with an active and partially-active state, respectively^8^.

The comparison of the resting and open or semi-open conformations shows that agonist binding promotes a contraction of the orthosteric site involving a large inward movement of the C-loop that caps the binding cavity. This is associated with a twist and tilt of each subunit β-sandwich. The tilt moves the upper part of the subunit ECD towards its neighbouring complementary subunit, while its lower part moves away from it. In consequence, the cys-loop at the bottom of the ECD undergoes an outward movement associated with TMD expansion, M2 tilt and channel opening. From this open-channel conformation, desensitization motions mainly concern the TMD that relaxes back towards an occluded, closed-channel conformation, associated with a subtle global expansion of the ECD.

In a previous study, we immunized alpacas with cells expressing an α7/5HT3 chimera to generate highly potent and α7-specific modulators^9^. Among the resulting camelid antibody fragments called nanobodies, the one called E3 acts as a Type I PAM. In two electrode voltage-clamp electrophysiology (TEVC), upon pre-application for 30-60 sec, E3 potentiates ACh-gated currents with a maximal effect around an EC_10_ concentration of ACh of 30 uM. E3 does not alter significantly the EC_50_ of ACh, nor the apparent slow desensitization component of the ACh-elicited response. In contrast, the nanobody called C4 acts as a silent allosteric modulator (SAM), showing no effect on ACh-gated currents but inhibiting E3-elicited potentiation. Both nanobodies do not displace Bgt in binding experiments, pointing to an allosteric binding site away from the orthosteric site.

In this paper, we aim at understanding the mechanism by which the nanobodies mediate their allosteric effect by solving the cryoEM structures of the complexes they form with the α7-nAChR. Our data enables the high-resolution identification of the common binding epitope targeted by C4 and E3, with in both cases an association of five nanobodies binding at the top of the receptor pentamer. Mutational investigations hint at a key N-linked sugar moiety in mediating E3 potentiation.

## Results

### 1/ Production of the purified human α7-nAChR reconstituted in nanodiscs

To express the α7-nAChR in mammalian cells, we used a lentivirus-based strategy^10^ (Figure S1). We designed two lentiviral constructs harbouring a gene coding for either the fulllength α7-nAChR subunit fused to a Rho1D4 tag at its C-terminus (α7FLcryo) or for the same gene where the coding sequence of the ICD is replaced by a short linker based on a α4β2 construct used for crystallogenesis^11^ (α7ΔICDcryo) (Figure S2). We additionally produced a lenviral construct harbouring the coding sequence of the chaperone NACHO^12^. Lentiviruses particles were used to infect T-REx cells grown in suspension. Expressed proteins were affinity purified in dodecyl-maltoside (DDM). As initial micrographs in detergent micelles showed protein aggregation, we further reconstituted the protein in MSP1E3D1 nanodiscs containing brain lipid extracts, resulting in well dispersed particles on cryoEM grids (Figure S1). Yields of production were improved around 2-fold in the α7ΔICDcryo construct as compared α7FLcryo. In parallel, we expressed and purified C4 and E3 nanobodies in a monomeric form as described previously^9^ (Figure S1 and S3)

We first verified that both α7cryo constructs retain the modulation properties of C4 and E3 in TEVC (Figures S1). As for the wild-type α7 nAChR, a pre-incubation of 1μM of E3 leads to the potentiation of ACh-gated currents with no apparent effect on desensitization. C4 pre-incubation has no significant effect by itself but inhibits E3-elicited potentiation (Figure S1). We also measured the binding affinity of both nanobodies for the nanodiscs-reconstituted α7ΔICDcryo using single cycle kinetics Surface Plasma Resonance assays. In both cases, Rmax values suggest a maximum of three nanobodies molecules bound per receptor molecule. Concentration-response curves show that C4 displays slower association and dissociation kinetics (k_on_ = 1.2±0.05 10^6^ M^−1^.s^−1^, ko_ff_ = 0.024 ±0.02 10^−2^ s^−1^) than E3 (k_on_ = 11 ±0.02 10^6^ M^−1^.s^−1^, k_off_ = 1.19 ±0.01 10^−2^ s^−1^). Data show high affinity for both C4 and E3 with K_d_ of 0.2 ±0.3nM and 1 ±0.01nM respectively (Figure S4). We also assayed E3 in the presence of nicotine, to investigate whether a possible increased affinity could account for its PAM action. However, the affinity of E3 was found identical in the absence and presence of nicotine (Figure S4).

### 2/ Cryo-EM structures of α7 in complex with C4 show preservation of nicotine-elicited reorganizations

We first solved the cryo-EM structure of the α7ΔICDcryo in the presence of a 50-fold excess of C4 (1μM α7 pentamer for 50μM C4, C4-Apo structure) (Figure 1 and S5, table S1). After 2D and 3D classifications, the final set of particles yielded a 5-fold symmetrical reconstruction, with five bound nanobodies per α7 pentamer. For most of the ECD and nanobodies, the density map is of good quality and allows to confidently build the main and side chains of the protein with a nominal resolution of 2.3 Å. In contrast, the density of the TMD and associated lipid nanodisc, although visible at low contouring of the map, shows poor quality and with no elongated and rod-like feature that could be attributed to transmembrane helices (Figure S7). The latch turn, a short helix protruding from the C-terminus of M4 and interacting with the lower part of the cys-loop, is not resolved. This shows that, in MSP1E3D1 nanodiscs containing brain lipid extracts, the TMD conformation of α7ΔICDcryo is heterogeneous among the selected particles, indicating structural flexibility.

In the published structures in detergent or nanodiscs^6,7^, the variable loop from the ICD was truncated but the two α-helices MA and MX were kept and resolved, notably the MA helices of each subunit that form a tight quaternary pentameric bundle that could possibly rigidify the TMD. To check if the weak TMD resolution is due to our ICD truncation, we solved the structure of α7FLcryo, also with a 50-fold excess of C4 and adding 100μM nicotine to further stabilize the protein conformation (C4-Nic structure) (Figure 1 and S5, table S1). The resulting reconstruction reaches a resolution of 3.6 Å for the ECD/nanobody part but also features a disordered transmembrane domain. Still, extra density is visible below the TMD/nanodisc area, compatible with the presence of the bundle of MA helices (Figure S7). In the previous cryo-EM structures of α7 obtained in saposin nanodiscs containing soy lipids/25% cholesterol, the TMD was built^6^. However, its local resolution was lower than that of the ECD, indicating that the flexibility of the TMD is a distinctive feature of the α7 nAChR.

Our data nevertheless allow the model building of the α7 ECD, including the first glycan linked to α7 at Asn23 and Asn67 and two glycans on α7 Asn110, in complex with C4. C4 adopts a classical immunoglobulin-like fold, despite lacking the canonical intra-sandwich disulfide bond (Figure S3). In both C4-Apo and C4-Nic structures, five nanobodies are bound in a similar way and form an additional ring above the ECD. We numbered the nanobody chains according to the α7 subunits anticlockwise arrangement, i.e. C4-chainF (*n*) sits above α7-chainA (principal subunit, *n*), C4-chain G (*n+1*) sits above α7-chain B (complementary subunit, *n+1*). Each nanobody interacts mainly with the top of a principal α7 subunit but also with the α7 complementary subunit, and presents its CDRs facing the ECD. The axis of each nanobody β-sandwich is roughly perpendicular to the membrane, in an orientation close to that of the β-sandwiches of the α7 ECDs, which also display an immunoglobulin-type fold. Nanobodies interact substantially with each other (see below), and, while monomeric in solution, they present here a 5-fold assembly. The detailed description of the receptor-nanobody and nanobody-nanobody interactions are provided in sections 4/ and 5/.

We then analysed the receptor global conformation by measuring deviations to known structures and looking at structural hallmarks conserved in the family (loop C capping, cys-loop motion). First, for the C4-Nic structure, Cα Root-mean-square deviations (rmsd) calculations of the α7 ECD shows that the C4-Nic structure aligns best with structures obtained in presence of agonists, with or without PNU (Figure 2 and S8, table S2). Superimposition of the core of the α7 β-sandwich nicely illustrates this point (Figure S8). Measurement of the Loop C capping (10Å for the Cys189_A_-Tyr117_E_ distance), and of the cys-loop outward motion (11Å for the Cys127_A_-Ile168_E_ distance) clearly shows values that are characteristic of active/desensitized rather than resting conformation (Loop C capping around 10Å vs 17 Å, cys-loop motion around 10Å vs 8Å, respectively).

In the five orthosteric sites of C4-Nic, we found an additional density in which we could confidently build a nicotine molecule (Figure 1 and 2). The pyrrolidine points downward, and its electropositive ammonium lies in a box made up of aromatic residues from loop A (Tyr92), B (Trp148), C (Tyr187, Tyr194) and D (n+1 Trp54). The pyridine points upward and elicits contacts with the loop C (disulfide bridge Cys189-Cys190) and loop E (n+1 Leu118). Similar poses of the agonist were found in the structure of heteromeric α4β2 and α3β4 nAChRs in complex with nicotine^11,13,14^, and in the α7 nAChR in complex with the related alkaloid epibatidine^6^. These observations are consistent with a conformation of α7 corresponding to an active, partially-active or desensitized conformation. Given that information about the TMD conformation is missing in our dataset, we cannot unambiguously distinguish between those possibilities.

**Figure 1 :**
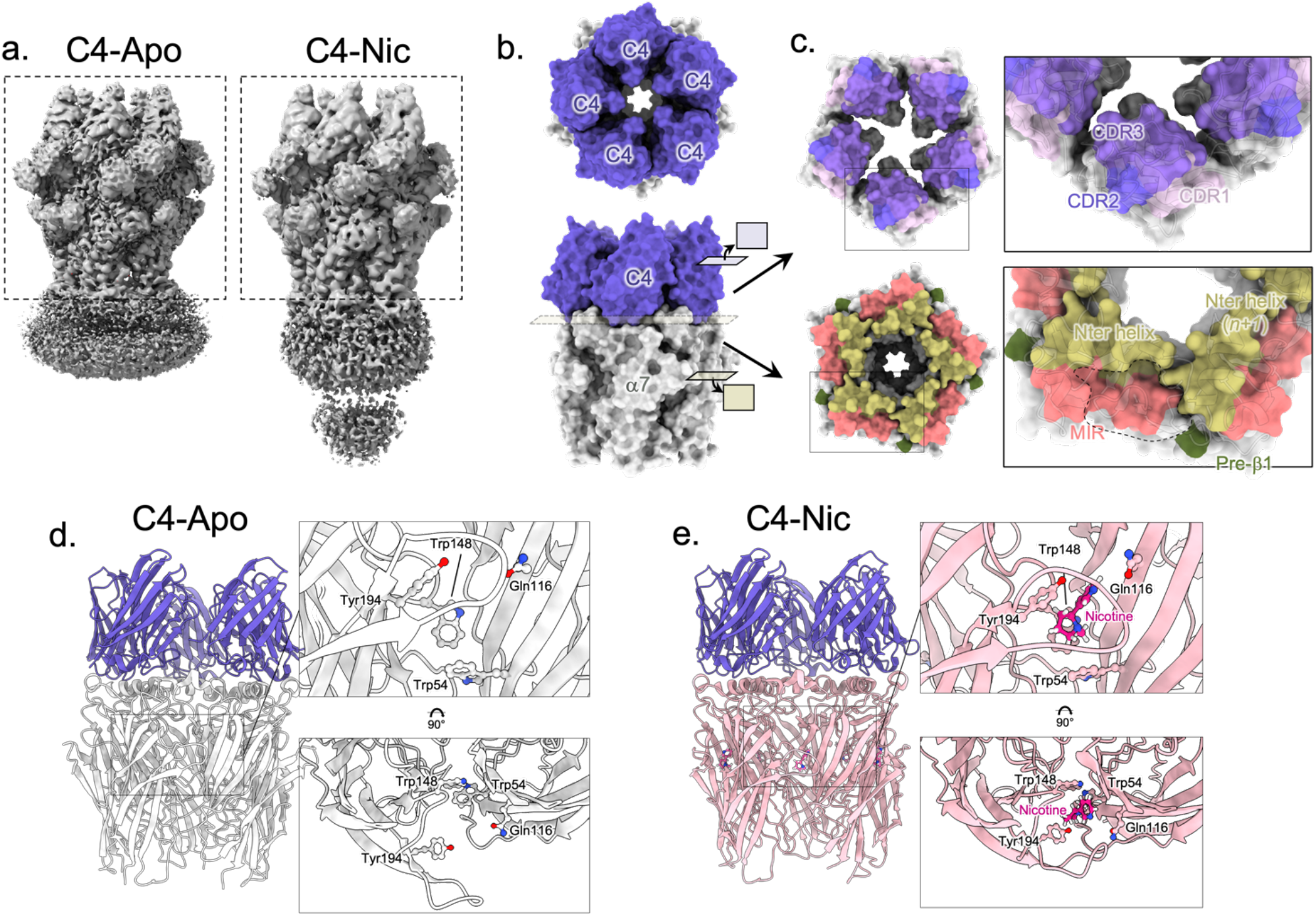
C4-α7 cryo-EM structures in the absence or presence of nicotine. a. Unsharpened maps of the α7ΔICDcryo in complex with C4 (C4-Apo, left) and of the α7FLcryo in complex with C4 and nicotine (C4-Nic, right) with densities contoured at 2σ. The dashed boxes denote the parts that were used for model building. b. Top and side views of the C4-Apo structure, protein chains are depicted in white (α7 apo-) and purple (C4). The structure is shown exploded in c. c. Top : Bottom view of the ring of C4 with the molecular surface of the three CDRs colored in shades of purple. Bottom : Top view of the ECD of α7 with Nter helices, MIR and pre-β 1 colored in yellow, pink and green respectively. The groove in which the CDRs are plunging is delimited with a dashed line. d. Side view of the C4-Apo structure, represented in cartoon with α7 in white and C4 in purple. The orthosteric binding site is enlarged and further rotated by 90°C in the right panels. Residues involved in nicotine binding are depicted in sticks. e. Side view of the C4-Nic structure, represented in cartoon with α7 in pink and C4 in purple. The orthosteric binding site is enlarged and further rotated by 90°C in the right panels. Residues involved in nicotine binding and nicotine itself are depicted in sticks.

**Figure 2 :**
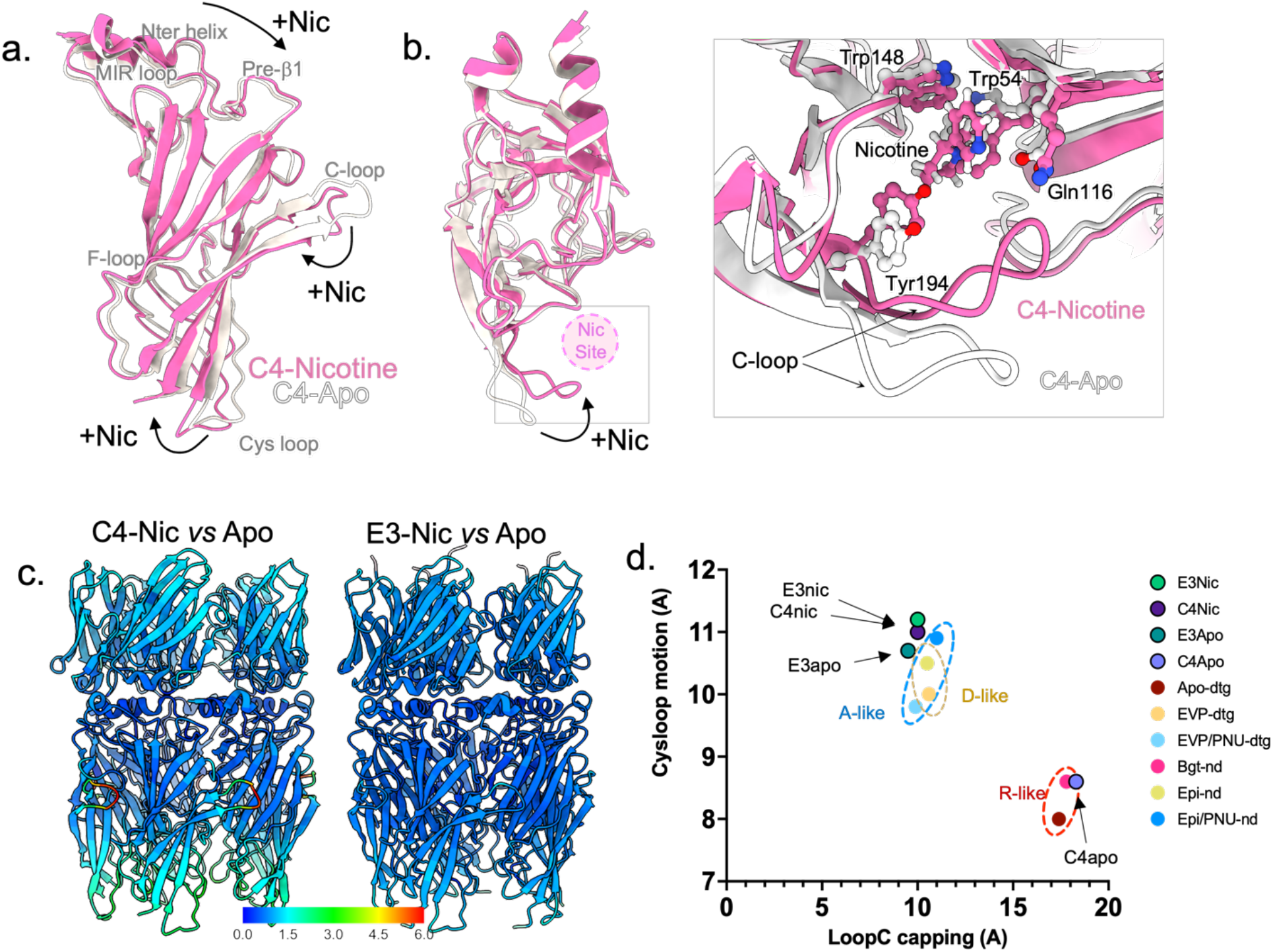
α7 conformation in C4-Apo corresponds to a resting-like state, and in C4-Nic, E3-Apo and E3-Nic to a partially active/active/desensitized state. a. α7 monomers from the C4-apo (grey) and C4-Nic structure (pink) are seen from the side and depicted in cartoons after superimposition of the whole α7 pentamer. Landmark loops are labelled and motions from Apo- to Nic- are highlighted with black arrows. b. The same representation than in a. but seen from the top. Nicotine binding site is depicted with a pink circle. A close view of the nicotine binding site is shown on the right. Nicotine and key residues from the binding pocket are labelled and depicted in sticks. c. Conformational changes upon nicotine binding in C4-bound (left) and E3-bound (right) structures. C4-Nic and E3-Nic are represented in cartoons and Cα are colored according to the rmsd value calculated using their Apo-counterpart, with the same range for both and shown in the color key below. d. Correlation of the cys-loop outward motion and Loop C capping in the known structures of α7 allows to define pairs of values that are landmarks of the α7 conformations. R, A, and D-like structures are grouped in dashed circles.

For the C4-Apo structure, the α7 ECD structure aligns best with that of the Bgt-nanodisc and apo-detergent structures (Figure 2 and S8), with notably an uncapped Loop C (17 Å for the Cys189_A_-Tyr117_E_ distance) and no cys-loop outward motion (Cys127_A_-Ile168_E_ distance below 9 Å). Those markers indicate that the C4-Apo structure shows a resting-like conformation of α7. This is best exemplified by the orthosteric binding site which is empty and displays a wide-open loop C (Figure 1). In the nanobodies region, comparison of C4-Apo and C4-Nic structures shows few conformational changes upon nicotine binding apart from a small motion of the apex of the nanobodies away from the pentameric axis (Figure S9).

In conclusion, with five C4 nanobodies bound to its ECD, α7 adopts conformations similar to those observed in the absence of nanobody, namely a resting-like conformation in the absence of agonist, and an active/partially-active/desensitized-like conformation in the presence of nicotine. Therefore, C4 appears to be a neutral binder, preserving the nicotine-elicited reorganizations in Cryo-EM conditions.

### 3/ E3 alone stabilizes an agonist-elicited conformation in Cryo-EM structures

We then solved the structure of α7ΔICDcryo with a 50-fold excess of the E3 nanobody, with and without 100μM nicotine, with an overall resolution of 2.6 and 2.8 Å in the ECD/nanobody region, respectively, and low signal in the TMD (Figure 3, S10, S11). Five E3 molecules are bound, in a similar arrangement to the one seen for C4. E3 adopts the same overall fold than C4 with local variations.

**Figure 3 :**
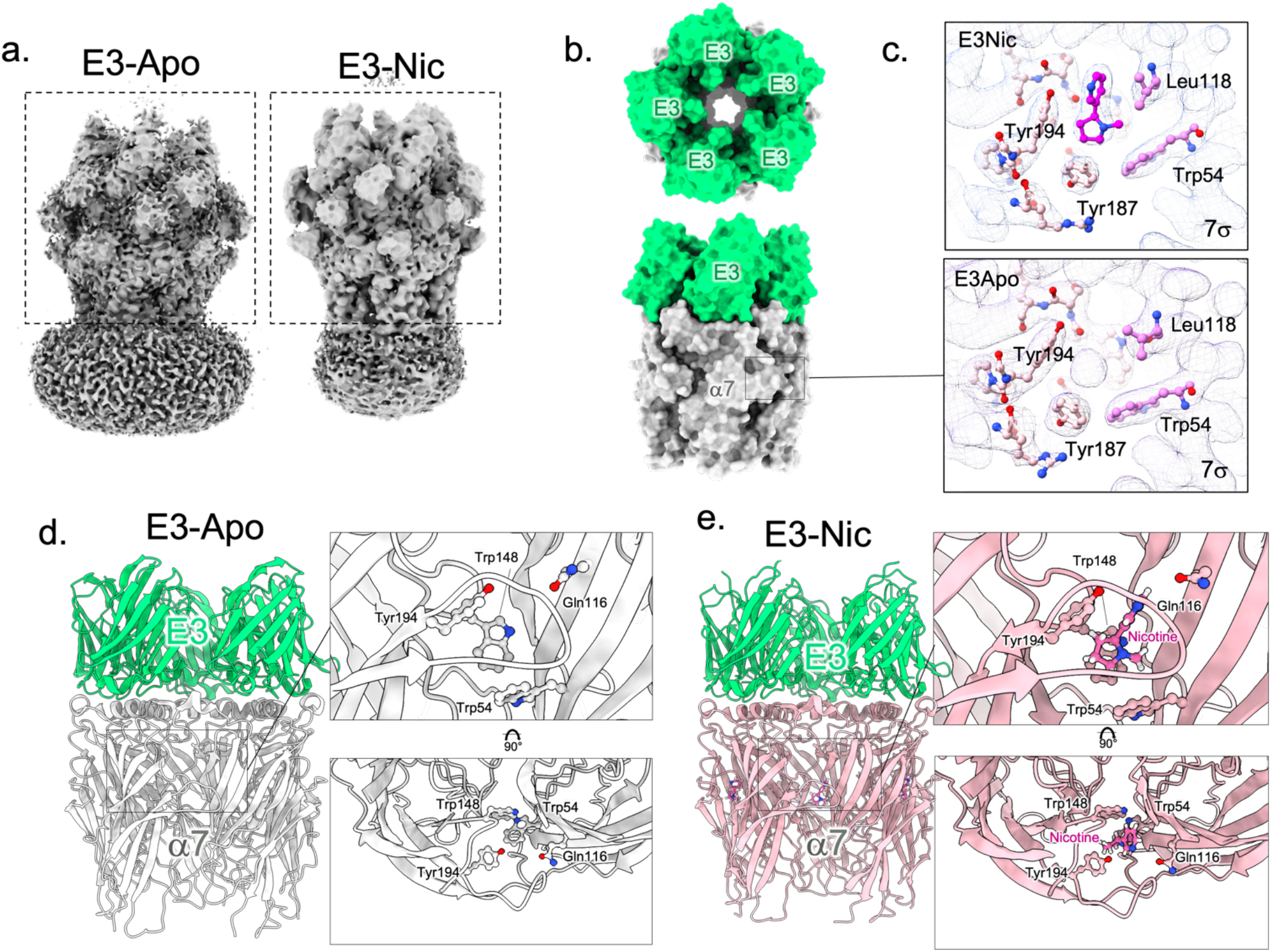
E3-α7 cryo-EM structures in the absence or presence of nicotine. a. Unsharpened maps of the α7ΔICDcryo in complex with E3-Apo (left) and E3-Nic (right) with densities contoured at 2σ. The dashed boxes denote the parts that were used for model building. b. Top and side views of the E3-Apo structure, protein chains are depicted in white (α7 apo-) and green (E3). The orthosteric sites are shown in c. c. Neurotransmitter binding pocket of E3-Nic and E3-Apo. Electron densities are contoured at 7σ and residues involved in neurotransmitter binding are shown in sticks. Nicotine was built in the extra density found in E3Nic while the small and spherical density found in E3-Apo likely fits a cation or a water molecule. d. Side view of the E3-Apo structure, represented in cartoon with α7 in white and E3 in green. The orthosteric binding site is enlarged and further rotated by 90°C in the right panels. Residues involved in nicotine binding are depicted in sticks. e. Side view of the E3-Nic structure, represented in cartoon with α7 in pink and E3 in green. The orthosteric binding site is enlarged and further rotated by 90°C in the right panels. Residues involved in nicotine binding and nicotine itself are depicted in sticks.

The E3-Nic structure shows an ECD conformation very close to that of C4-Nic, including the five bound nicotine molecules, consistent with active/partially-active/desensitized conformations (Figure 2, 3, S8 and S9). Surprisingly, the conformation of the α7 ECD in the E3-Apo structure also matches best the agonist-bound conformations, including Loop C capping and Cys-loop outward motion (Figure 2 and 3). At the orthosteric site, the conformations of all binding determinants resemble the ones seen in the E3-Nic structure. In the cavity, we found a well-defined small spherical density, which is located where the electropositive pyrrolidine ammonium would be. The density would be compatible with water, or more likely a cation such as Na^+^ that may contribute, along with the binding of E3, to compact the orthosteric site in the absence of nicotine (Figure 3). Yet, Cα rmsd calculations show that E3-Apo slightly differs from both E3-Nic and C4-Nic, with locally 1-2 Å deviations in the loop C, as well as the top and bottom regions of the ECD (Figure S9). In addition, we could observe a slight expansion of the ECD subunit in E3-Nic vs E3-Apo (~1 Å displacement), a motion comparable with the one observed between the Epi vs Epi/PNU structures in nanodiscs (PDB 7KOQ and 7KOX, Figure S9) but absent between the EVP and EVP/PNU structures in detergent (PDB 7EKP and 7EKT). As seen with C4, nicotine binding triggers a small motion of the tip of the nanobody away from the channel axis (Figure S9).

In conclusion, our data point to a striking effect of E3, which, in the cryo-EM conditions (high 50μM E3 concentration, 1 hour equilibration period before freezing in the grids) stabilizes by itself an agonist-bound conformation through long-range allosteric interactions initiated at the top of the receptor. This suggests that, in electrophysiology, high E3 concentrations and long incubation periods could also trigger activation and/or desensitization in the absence of orthosteric agonist. However, such conditions are hardly reachable in a TEVC experiment. In addition, the fast desensitization of α7 could mask a slow gradual activation by E3. To overcome these issues, we used the α7 mutant L247T that disrupts a central hydrophobic ring in the channel^15,16^ (Figure 6). This mutant strongly reduces desensitization and produces a marked gain of function phenotype increasing the efficacy of partial agonists. We recorded L247T currents elicited by 0.3μM ACh before and after a 10 sec pre-application of 10 μM of E3. As expected, ACh-gated currents are potentiated by E3 pre-application (2.4 ±1.1 fold). In addition, we observed an increase of the holding current during the pre-application of E3 that represents 6.0 ±1.8 % of the response to 0.3μM ACh. This shows that E3 does have significant agonist properties in the absence of ACh when applied for a short period at a relatively high concentration on a weakly-desensitizing mutant.

### 4/ Analysis of the E3 and C4 common epitope

C4 and E3 bind to α7 in a highly similar manner (Figure 4 and S4), their CDRs contacting the top-ECD platform formed by the principal α7 subunit down below (with a ~500 Å^2^ surface of interaction), and the complementary α7 subunit (~300 Å^2^). On the principal subunit, it is formed by the C-terminal half of the top α helix that faces the vestibule, and by the long β2-β3 loop, also called Main Immunogenic Region (MIR)^17^, on the outer side. On the complementary subunit, it is formed by the N-terminal half of the top α-helix.

**Figure 4:**
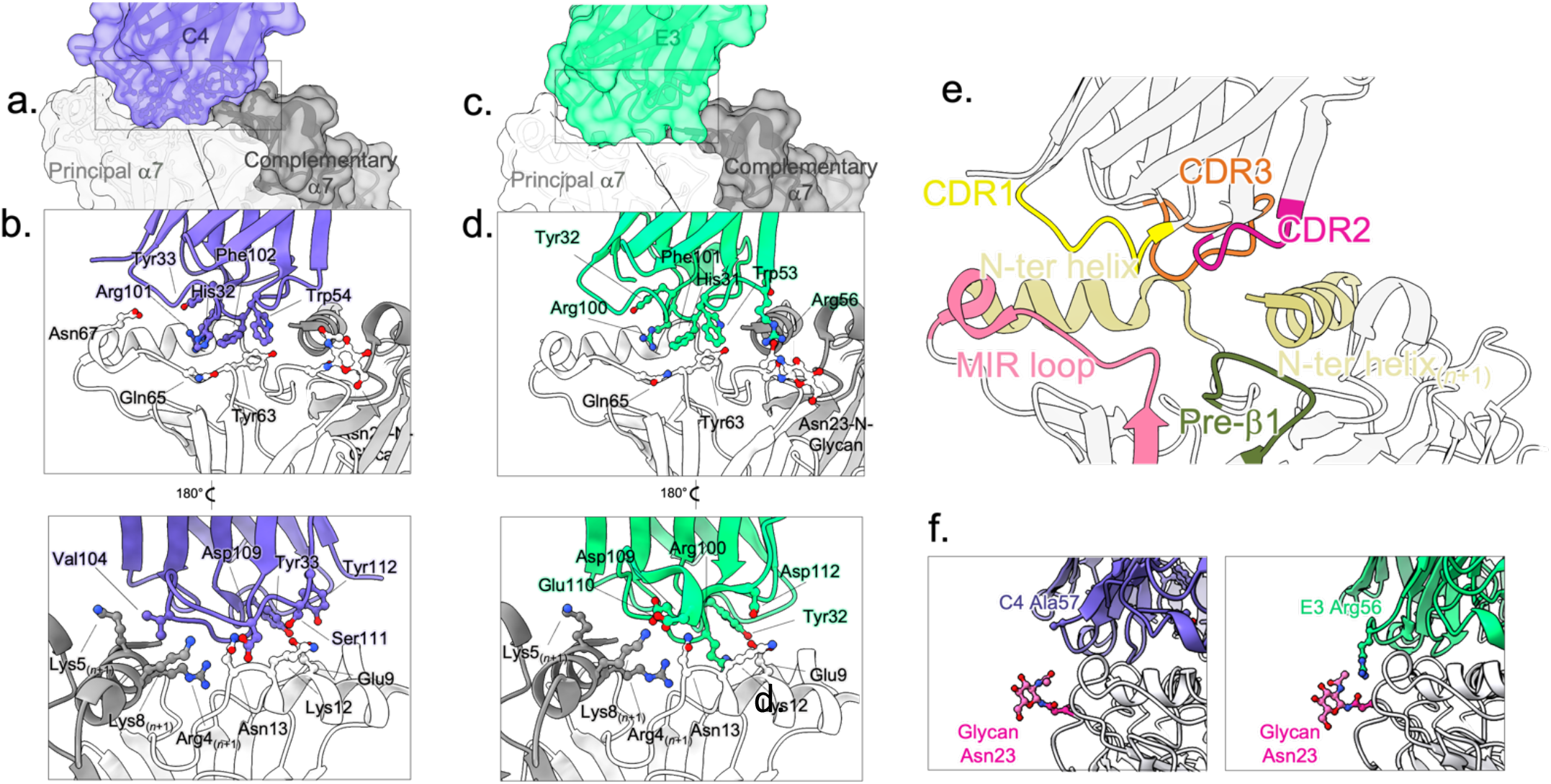
Binding pose and epitope of C4 and E3. a. Overview of the binding site of C4 on α7 exemplified by the C4-Apo structure. One C4 molecule (purple surface and cartoon) contacts two α7 subunits depicted in light (principal n subunit) and dark (complementary anticlockwise n+1 subunit) surface and cartoons. b. Details of the binding site as seen from the solvent (upper panel) or the vestibule (lower panel) of α7 with color code as in a. key residues are labeled, and their side chains displayed in stick balls. c. Same as in a. but with the E3-Apo structure, E3 is colored in green d. Same as in b. but with the E3-Apo structure, E3 is colored in green. e. Overview of the loops involved in binding exemplified by the C4-Apo structure represented in cartoons. f. Interaction of E3 with Asn23 glycan. Close view of the Glycan (pink sticks) built on α7-Asn23 (pink sticks) from the pre-β1 of α7 (grey) and of the CDR2 of C4 (left, purple) or E3 (right, green). Ala57 from C4 and its homologous Arg56 from E3 are depicted in sticks.

Both nanobodies display unique disulfide bond patterns. First, E3 harbours the “canonical” disulfide bond that links the strands that just precedes the CDR1 and CDR3, which is missing in C4. Second, the CDR3 of E3 is longer by two residues, and is bent upwards by the presence of an extra disulfide bond between the end of the CDR3 and the second β-strand of the framework region 3 (FR3) (Figure S3). In known nanobodies featuring a long CDR3, a supplementary disulfide bond rigidifying the structure is common^18^. For both nanobodies, conserved side chains from the three CDRs (His32_CDR1_, Trp54_CDR2_, Arg101_CDR3_ and Phe102_CDR3_, C4 numbering) are directly plunging into a groove bordered by the helix and MIR elements. Both nanobodies elicit a complex pattern of interactions with α7 that is detailed in tables S3 and S4. Although the sets of interactions are different in C3 and E4, they involve conserved residues from the CDR1, CDR2 and first half of CDR3 (Gly27_CDR1_, Tyr33_CDR1_, Trp54_CDR2_, Arg101_CDR3_, Phe102 _CDR3_, C4 numbering). They involve also non-conserved residues from the second half of the CDR3, where C4 residues Asp109 and Ser111 are engaged in an extensive set of interactions with the principal N-terminal helix, while much fewer contacts are found in E3 (Figure 4, tables S3 and S4). In addition, E3 shows a unique interaction between the Arg56_CDR2_ guanidinium and both the main chain carbonyl and the first glycan grafted on α7Asn23 (pre-β1), an interaction absent in C4 that harbours an alanine in place of the arginine at this position (Figure 3). Overall, the CDR3 is the main interacting loop, contacting the MIR and the N-ter helices of both subunits.

We further investigated the contribution of specific α7 motifs to nanobody binding by mutagenesis of α7FLcryo. On the one hand, we mutated the charged residues of the Nter helix of α7 (R4Q K5Q K9Q triple mutant and E9Q K12Q N13A triple mutant), and on the other hand we replaced the MIR residues to the aligned MIR of the α3-nAChR which does not bind C4 and E3^9^. Finally, we prevented N-linked glycosylation by mutating the serine of the Asn-X-Ser motif (Ser25Ala). We performed immunofluorescence assays in non-permeabilized HEK293 cells (Figure S4). Immunolabelling of the extracellular C-terminal Rho1D4 tag show that the MIR and glycosylation mutants are expressed at the cell surface, while mutants of the Nter helix are not. Immunolabelling with E3 and C4, using nanobodies fused to the heavy chain fragment of a human IgG (constructs detailed in^9^, Figure S4), show clear labelling of cells expressing the WT and glycosylation mutant, but no labelling of the MIR mutant. This confirms that the MIR is essential for the nanobody binding, while the glycosylation on Asn23 is not mandatory for E3 high-affinity binding.

### 5/ Inter-nanobody interactions

A striking feature of the structures of α7 bound to C4 and E3 is the 5-fold assembly adopted by the nanobodies above the receptors (Figures 1 and 3), where the FR3 region of each C4/E3 interacts with the FR1 region of the *n+1* C4/E3 nanobody through a ~200 Å^2^ interface (Figure S8). In C4, Tyr60, Ser63 and Lys 66 contact n+1 Gln2 and Gln4 (Figure 5) and in E3, Asn62 and Lys65 contact n+1 Gln3 and Gln5 and Gln 116. The CDR3 of E3 also contributes to the interface, the guanidinium of Arg108 lying near the carboxylate of Asp 113 suggesting a salt-bridge interaction. A more extensive set of inter-nanobody interactions is thus observed in E3 vs C4.

**Figure 5:**
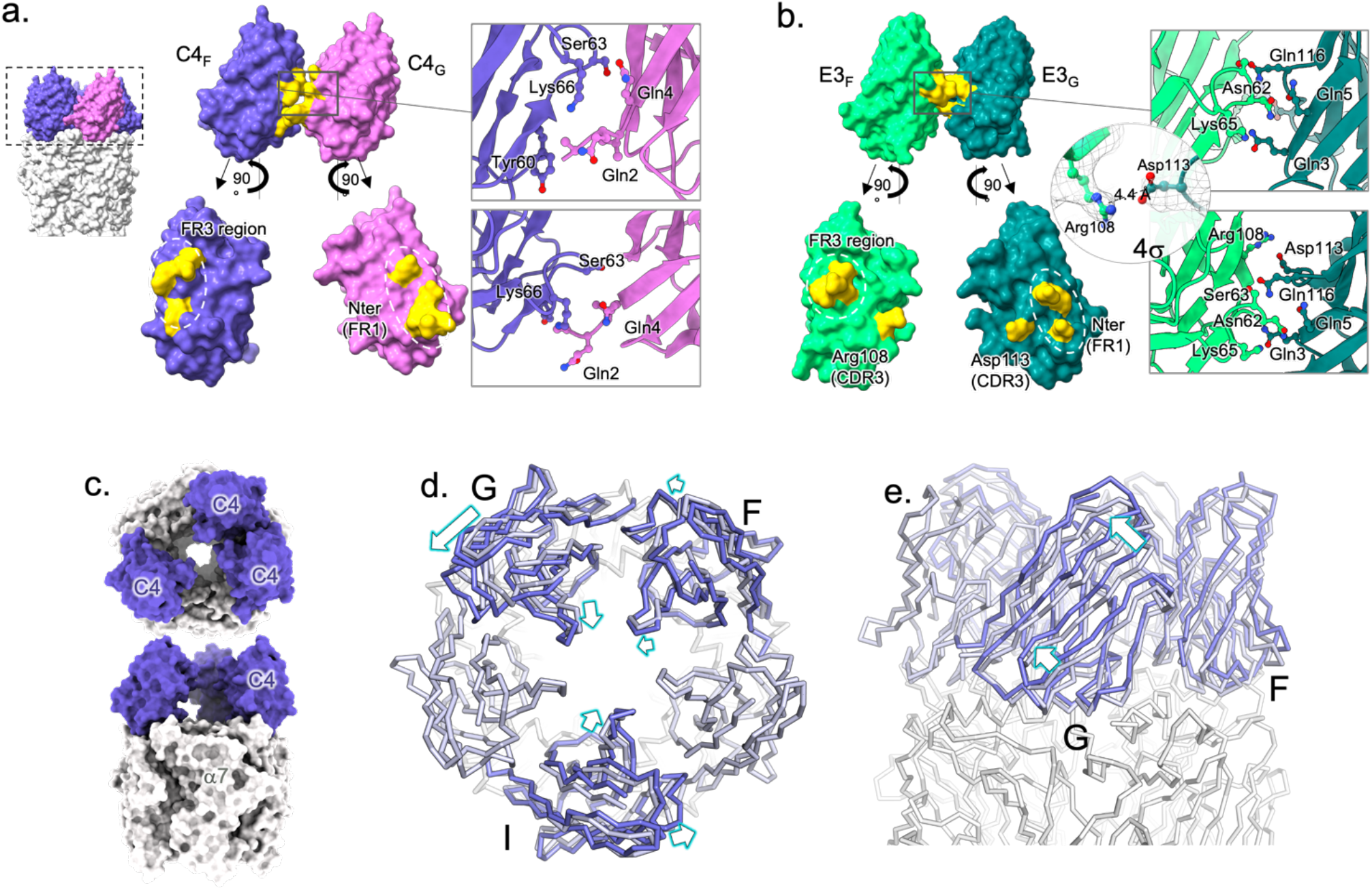
Nanobody-nanobody interactions. a. Surface representation of two adjacent C4 (F in purple, G in pink) in C4-Apo and their exploded view below. Residues at the interface are colored in yellow and their details are shown in cartoons and sticks in a close-up view (right). b. Surface representation of two adjacent E3 (F in light green, G in dark pink) in E3-Apo and their exploded view below. Residues at the interface are colored in yellow and their details are shown in cartoons and sticks in a close-up view (right). Additionally, the putative salt bond between Arg108 and Asp113 is detailed in a separate close-up view, with the density contoured at 4σ. c. Overview of the C4partial-Apo structure from the top (top) and the side (bottom) with α7 in grey and the three C4 molecules in purple. d. Superimposition of the C4partial-Apo on the C4-Apo structures, aligned on α7 ECD. Both structures are represented in ribbon, α7 in grey, C4 from C4partial-Apo in purple and C4 from C4-Apo in grey blue. Motion from C4-Apo to C4partial-Apo are highlighted with arrows. On the left, structures are seen from the top and show a lateral inclination of the nanobodies toward the anticlockwise subunit accompanied by a radial inclination towards the vestibule. On the right, structures are seen for the side, with the most displaced nanobody in front (chain G).

To further explore the quaternary binding mode of the nanobodies, we solved the structure of α7ΔICDcryo in the presence of a sub-saturating concentration of C4 (10μM) (Figure 5 and S11; Table S1). Indeed, in the early stages of the cryoEM work at the 2D classification step, we noticed that, for both E3 and C4, addition of 10μM of nanobodies is not sufficient to reach the C5-symmetry in the nanobody regions, indicating partial nanobody binding (Figure S13). We processed a dataset obtained with 10μM of C4 and no nicotine. 2D class averages show a minor fraction of particles containing one or two bound nanobodies, and a major fraction containing three bound nanobodies. We selected these later particles to solve a C4partial-Apo structure. Without imposed symmetry, we obtained a final volume of 4.4 Å resolution and, after density modification in Phenix^19^, we built a model with an overall model-versus-map resolution of 3.4 Å, where the main chains and most of the side chains of the α7 ECD and the three nanobody molecules are confidently built. For simplicity, we labelled the three C4 according to their position in C4-Apo: C4 chains F, G and I sit above α7 chains A, B and D, respectively. Interaction determinants of the three C4s with α7 are identical to the ones observe in C4-Apo. C4_F_ and C4_G_ interact with each other in a similar manner than in C4-Apo, although through a slightly larger interaction surface (~260 Å^2^, Figure S8), while C4_I_ sits alone.

Interestingly, each C4 chain displays a different radial and lateral inclination. The F, G and I chains are more inclined toward the vestibule (radial angles of 129.3°, 128.8° and 126.0°, respectively), compared to C4-Apo and C4-Nic (133.4° and 133.8°) (Figure S13). Likewise, the F, G and I chains are more inclined toward the (n+1) adjacent subunit (lateral angles of 82.8°, 80.5° and 78.2°, respectively), compared to C4-Apo and C4-Nic (85.1° and 85.3°). Therefore, data show that the nanobodies can bind independently from each other but that their orientation is constrained by the presence of neighbouring nanobodies. This suggests a sequential binding mode, where a single bound nanobody adopts a marked inclination toward the vestibule and the (n+1) adjacent α7 subunit, where two adjacent nanobodies show intermediate inclinations, and five bound nanobodies adopt a straighter binding orientation. This later is necessary for the α7 receptor to accommodate five bound nanobodies without steric clashes, eventually generating a symmetrical bundle (Figure 5).

### 6/ E3-elicited potentiation involves interaction with Asn23-linked glycan

The Cryo-EM structures show a sticking allosteric action of E3, a feature that parallels its PAM activity in electrophysiological experiments. In contrast, C4 appears “silent” with both techniques. We thus investigated whether specific E3-α7 contacts are involved the PAM activity. To his aim, we compared the E3 and C4 co-structures, searching for specific motifs/interactions of E3 absent in C4, followed by mutational and functional assays.

First, to investigate a possible contribution of the CDR3 conformation of E3, we removed the extra disulfide bond in the C95S/C114V E3 mutant, but its displays unaltered potentiating effect in TEVC (Figure 6).

**Figure 6:**
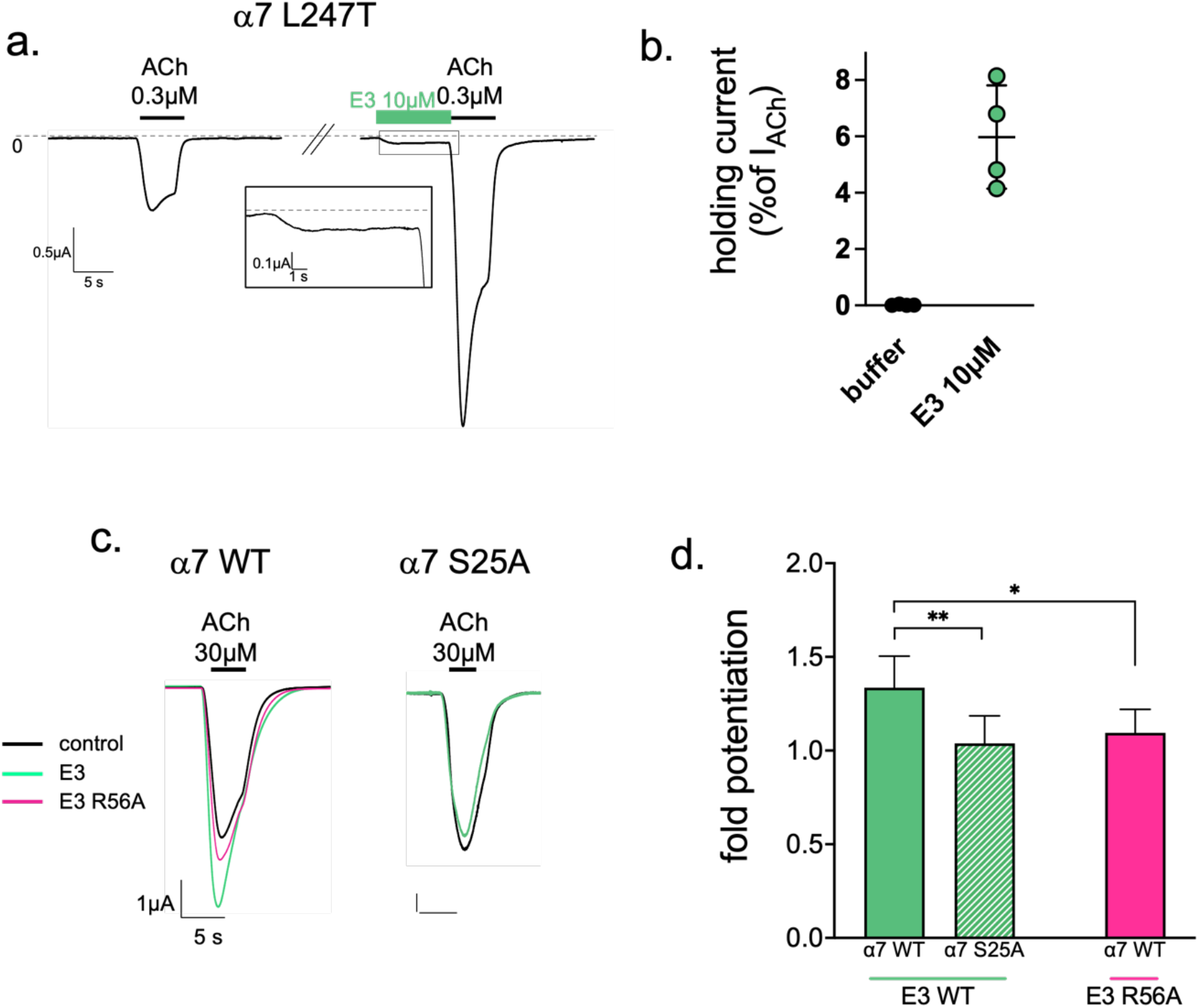
Electrophysiological analysis of the potentiation by E3. a. E3 direct activation on α7 L247T mutant recorded by TEVC on *Xenopus* oocytes. Representative trace showing a first 0.3μM ACh application yielding slow-desensitizing currents. After 2-min washing, 10μM E3 was applied for 10 sec, evoking significant current, before 0.3μM ACh. A closeup view of the recording during E3 application is shown in a black box. b. Holding currents measured on α7 L247T with or without 10 μM E3 normalized with the nonpotentiated response to 0.3μM ACh. c. Potentiation assays of E3-WT and E3-R56A on α7 WT and of E3-WT on α7-S25A. Responses to 30 μM ACh in the absence of E3 (black) are superimposed with response after 30 sec application of 1μM of E3 wild-type (green), or R56A (pink). d. Fold potentiation calculated on n≥4 cells by E3-WT on α7 WT and S25A mutant and of E3-R56A on α7-WT. Values were submitted to a unpaired t-test with * p≤0.05, **p≤0.01

**Figure 7 :**
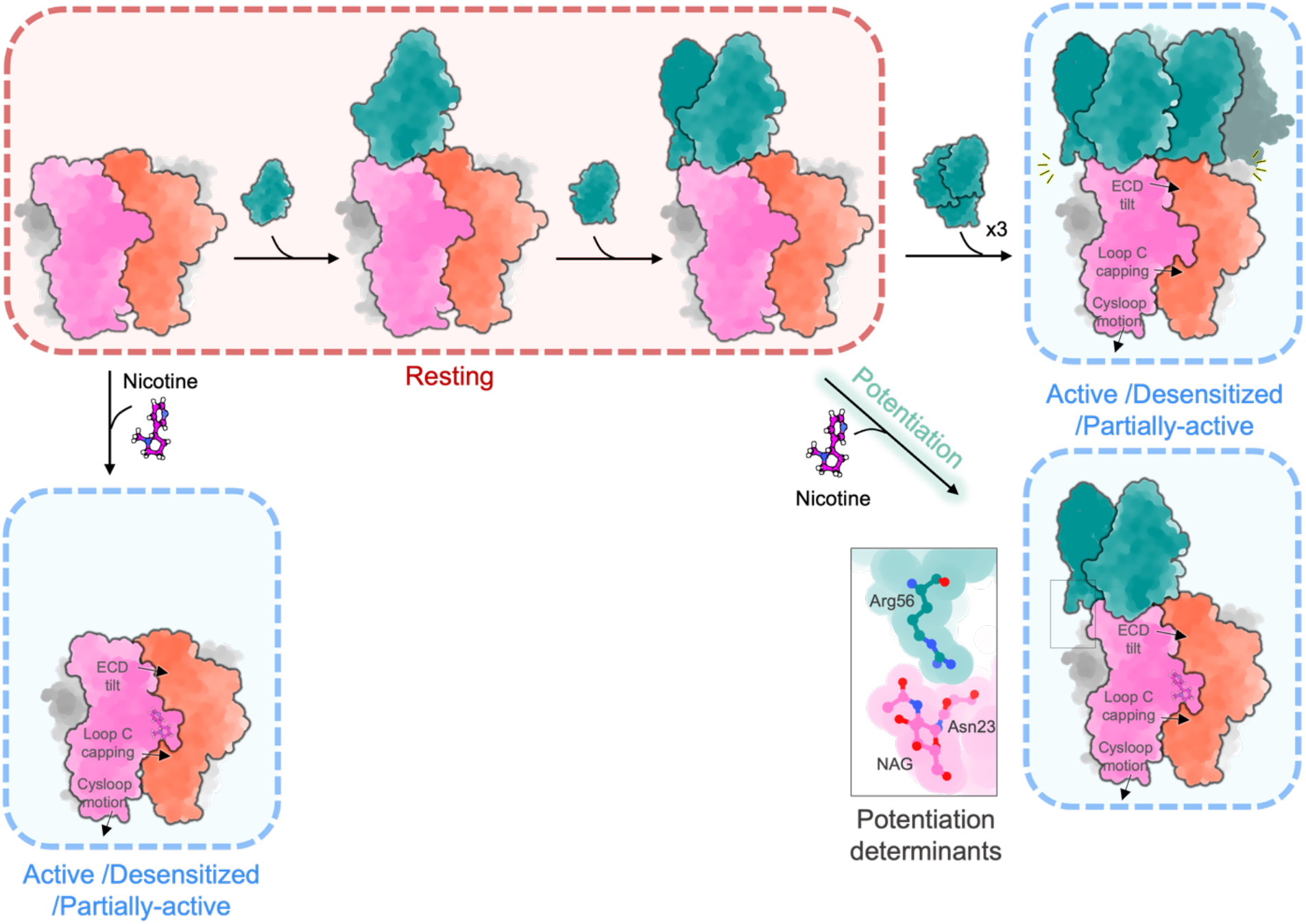
Hypothetical model for the allosteric mechanism mediating the PAM/agonist activity of E3. Cartoon representation of E3 (green) in complex with the α7 ECD (with two subunits colored pink and orange). Nicotine alone is shown to elicit the transition for the resting to the active/desensitized conformation. This transition involves ECD tilt, loop C capping and cys-loop outward motion. Our data suggest that, in typical TEVC conditions with an E3 concentration bellow 1 μM, potentiation of ACh-elicited response is caused by partial occupancy of the E3 binding sites, as arbitrary illustrated here by the binding of two nanobodies. Interaction between Arg56 of E3 and a N-linked sugar of α7 nearby the α7-α7 interface is critical for this potentiation. Cryo-EM data further show that higher concentrations (50 μM) of E3 are required to saturate the receptor, a condition in which E3 triggers the activation/desensitization conformational change in the absence of nicotine.

Second, a key feature of both nanobodies is their binding to the n+1 nanobody and α7 subunit. Since α7 undergoes a substantial quaternary reorganization during gating, the interaction of E3 with the complementary chains could contribute to constrain the relative arrangement of two adjacent α7 subunits toward an active/desensitized conformation. We mutated specific potential salt bridges between E3 and E3(n+1) (Arg108-Asp113), as well as between E3 and α7(n+1) (Glu110-Lys8_α7*n+1*_). E3-R108Q and E3-E110Q show an intact affinity by SPR, and a slightly weaker, yet not significantly different from E3, PAM activity by TEVC (Figure 6 and S4).

Third, a unique feature of E3 is the interaction between Arg56_CDR2_ and the first sugar moiety linked to Asn23 from the principal subunit. We showed in mutating α7S25A that the glycosylation is not mandatory for E3 high-affinity binding. Interestingly, α7S25A is no more potentiated by E3, suggesting a key contribution of the first sugar moiety in potentiation. However, we cannot exclude that the mutation, although qualitatively showing binding in immunofluorescence, could decrease the affinity for E3 to such an extent that weak binding would occur in our electrophysiological assay (1μM E3). To address this issue, we tested the E3-R56A mutant. It shows an affinity identical to that of E3-WT in SPR (Figure S4), and almost totally loses its ability to potentiate ACh-gated currents in electrophysiology (Figure 6 and S4).

In conclusion, data pinpoint an Arg-sugar interaction at the E3-α7(n) interface which mutation strongly decreases the potentiation, and two putative salt-bridges at the E3-α7(n+1) and E3-E3 interfaces whose mutation show a tendency to decrease the potentiation.

## Discussion

In this study we solved five structures of the α7-nAChR in complex with two allosteric nanobodies, unravelling a common epitope at the apex of the receptor which is formed by the MIR and the N-terminal helices from two adjacent subunits. In the pLGIC family, nanobodies were previously generated and used mainly to assist structural determination. Most bind at the ortho- or pseudo-ortho-steric site in the 5HT_3_R^20^, GABA_A_R^21,22^ and prokaryotic homolog ELIC^23^, except for a single NAM nanobody binding in the vestibule of ELIC^23^. It is noteworthy that E3 and C4 were generated through immunization of alpacas with cells expressing the fully glycosylated α7-nAChR ECD, while previous nanobodies involved immunization with purified proteins partly depleted from N-linked glycosylation by enzymatic cleavage (5HT_3_R) or production in GntI-cells (GABA_A_R). Interestingly, the unsharpened map of the E3/C4- bound structures suggests that glycosylation trees could substantially mask the orthosteric site, thereby plausibly preventing the generation of binders targeting this site and favouring a more apical epitope. Of note, the epitope of C4 and E3 shows weak sequence conservation among the various nAChR subtypes, accounting for their high specificity to the α7-nAChR with no observed binding by immunofluorescence to the α4β2 and α3β4 nAChRs^9^.

Several Fab fragments of conventional IgG were also found to bind within or around the orthosteric and pseudo-orthosteric sites in GABA_A_R^24,25^ and α4β2 nAChR^13^. More importantly, the MIR was first identified in the α1 subunit of the muscle-type nAChR as a key epitope for autoantibodies causing *myasthenia gravis*^17,26^. The crystal structure of the monomeric ECD of α1 in complex with a Fab fragment of one of those autoantibodies shows a binding mode involving interaction of both Fab chains with the MIR and the N-terminal helix of α1^27^. A somewhat similar binding mode is seen for an anti-α3 Fab in complex with the α3β4 nAChR^14^. However, both structures show a pose of the antibodies tilted to the edge of the top-ECD platform, the epitope centred on the MIR rather than the groove formed between the Nter helices and the MIR, as seen with E3 and C4. Therefore, the epitope of E3 and C4, while containing the MIR region, is unique among those so far characterized in pLGICs.

A unique feature of C4- and E3-bound structures is also the “pentamerization” of the nanobodies when binding to the α7 template. The C4partial-Apo structure shows that C4 binds in a sequential manner, a process requiring significant reorganization (change of radial and lateral inclination) of already bound nanobodies to accommodate new ones. Interestingly, SPR shows that C4 displays 0.1nM affinity with an evaluated maximal occupation of 3 nanobodies per pentamer. In contrast, Cryo-EM experiments show that 10 μM of C4 is not sufficient to saturate the receptor, full saturation requiring a 50 μM concentration. Data thus suggest that, within α7 pentamers, the first binding events display high affinity (nanomolar from SPR) while the last binding events would display micromolar affinity. We suggest that the sequential binding mode, where the binding of nanobodies is impaired when adjacent positions are already occupied, might cause this negative cooperativity. This mechanism might also apply to E3 which displays similar difference in SPR vs Cryo-EM nanobody binding features.

Cryo-EM and TEVC experiments provide compelling evidence for a strong allosteric effect of E3, but not of C4. In this line, removal of the sugar linked to Asn-23, that contributes to the E3 but not to the C4 epitope, strongly impairs the PAM activity of E3 by TEVC. This indicates that the binding of E3 to the top platform drives both 1/ the potentiation of ACh-elicited currents on the WT α7 and 2/ the stabilization of the active/desensitized state in the absence of orthosteric agonist, as seen in Cryo-EM and on the α7-L247T mutant. The PAM activity of E3, evaluated by potentiation of current elicited by 30 μM of ACh, is maximal above 1 μM with an apparent EC_50_ around 100 nM. SPR measure an affinity of 1 nM, suggesting that several binding events might be required to detect a PAM activity. In contrast, low μM concentrations of E3 are not sufficient to detect a direct agonist action. We show that 10 μM E3 alone weakly but significantly activates the α7L247T mutant, and that 50 μM E3 alone drives α7 into an active/partially-active/desensitized conformation in cryo-grids. Direct action of E3 is thus likely elicited by near-complete or complete saturation of the binding sites on the α7 pentamer.

E3 mediates allosteric effects through its binding at the top platform of the receptor. Beside the orthosteric site, the α7 nAChR is known to carry allosteric sites at the TMD, binding notably ivermectin^28^ and PNU^5,7^, at the ECD-TMD interface where calcium binds^29^, and at a vestibular site located nearby the orthosteric site, that binds small fragments^30^. In the pLGIC family, another allosteric site is well established, the binding site of AM-3607, a PAM binding between the top platform and the orthosteric site of the α3GlyR^31^. Thus, the allosteric site targeted by E3 is unique and involves a large surface involving multiple contacts. E3 is strongly anchored in the principal α7 subunit down below, with bulky side chains plunging into a groove bordered by the MIR and the helix. The PAM activity is strongly impaired by disrupting a key polar interaction with a sugar from the principal subunit nearby the interface with the complementary subunit, and possibly two putative salt bridges with the complementary n+1 E3 and α7 subunit. We speculate that the binding of E3 constrains the relative arrangement of the principal and complementary subunits of α7, favouring a subunit tilt associated with gating (Figure 7). Therefore, E3 implements a novel allosteric site which is the most remote from the core of the coupling pathway between the orthosteric site and the activation gate of the channel.

In conclusion, our work reveals a unique mechanism where nanobodies prolong the length of the pentameric architecture of the ECD to control the global allosteric conformation of the receptor. The modular architecture of pLGICs, made up of ECD and TMD domains that fold in a partially autonomous way^32–34^, is thus artificially extended here by an extra-module that tightly binds and is allosterically coupled to the native receptor. Such a regulation by additional modules is reminiscent of receptors carrying a regulatory N-terminal domain such as the pLGIC Declic^35^, or the tetrameric glutamate receptors. Finally, from a pharmacological perspective, E3 constitutes an original and highly subtype-specific therapeutic antibody candidate for α7-linked pathologies, including autoimmune^36^, neurodegenerative or psychiatric diseases.

## Methods

### Lentiviral vectors construction and production

The lentiviral transfer plasmid was generated from the pHR-CMV-TetO2-IRES-EmGFP and pHR-CMV-TetO2-IRES-mRuby2 plasmids available on Addgene. Human cDNA for α7-nAChRs and NACHO inserts were cloned using In-fusion cloning kit (Takara). After cloning in the lentiviral transfer plasmids, genes are preceded by a CMV promoter and a tet repressor element (CMV-TetO2) and followed by an IRES element promoting the expression of GFP (IRES-GFP, α7) or mRuby (IRES-Ruby, Nacho).

Viral particles were generated by co-transfection of HEK-293T cells by the lentiviral transfer plasmid, a packaging plasmid and an envelope plasmid. Two days after transfection, viral particles were harvested in the supernatant, treated with DNaseI, filtered through 0.45μm pores, concentrated by ultracentrifugation and resuspended in PBS. Viral stocks were stored in small aliquots at −80°C before use. Viral titres were estimated by quantification of the p24 capsid protein using HIV-1 p24 antigen immunoassay (ZeptoMetrix) and by Fluorescence Activated Cell Sorting (FACS).

### Lentiviral transduction, inducible expression in Trex cells

A mixture of α7-containing and NACHO-containing lentiviral particles (10:1) was used to transduce HEK T-Rex 293 cells (ThermoFischer). The resulting inducible cell line was cultured in suspension flasks in Freestyle 293 expression medium supplemented with blasticidin (5-10μg/ml final) and FBS (1%) in an orbital incubator at 37°C and 8% CO2. Protein expression was induced at a density of 3-4×10^6^ cells /ml by addition of doxycycline (10μg/L final). The cells were further cultured for 48h before harvesting by centrifugation.

### α7-nAChR purification

After low-speed centrifugation, the pellet was washed with cold PBS 1x and centrifuged again. Cells were resuspended in buffer A (20mM Tris pH8.0, 250mM NaCl) supplemented with antiprotease cocktail and mechanically lysed (Ultra-turrax T20, IKA, 9×30 seconds). Lysed cells were centrifuged at low speed (2000g) to remove debris and intact cells and membranes were collected by ultracentriguation (190000g for 2 hours). All further steps were carried out at 4°C. Membranes were mechanically broken and were resuspended in buffer A (10ml of buffer per gram of membrane) and supplemented with 1% w/v final of dodecyl-maltoside (DDM, Anatrace) for 1h solubilization under gentle stirring. The insoluble material was removed by ultracentrifugation (190000g for 45 minutes). The supernatant containing solubilized proteins was bound overnight at 4°C on a gravity flow Rho1D4 resin (Cube Biotech) equilibrated with buffer A supplemented with 0.05% DDM. After binding overnight, resin was washed with buffer A and 0.05% DDM and gradually washed to remove DDM and exchangeit for 0.01 % Lauryl Maltose Neopentyl Glycol (LMNG, Anatrace).

MSP1E3D1-Histag plasmid was obtained from Addgene, and protein was purified as described in Ritchie *et al* 2009. Porcine Brain Lipids Total extracts (BLT, Avanti) were resuspended in buffer A containing 0.01% LMNG at 10mM, sonicated and stored at −80°C until use.

The α7 receptor on its resin was incubated with MSP1E3D1 and BLT extracts (ratio 1:2:200) for 1h at 4°C. The removal of detergent was initiated by the addition of bio-beads (BioRad) twice for 1h at 4°C. The receptor-lipidic disc mixture was washed in buffer without DDM and was eluted with 500μM Rho1D4 peptide (Cubebiotech) for 4×1hour at 4°C. The receptor-lipidic disc mixture was concentrated to 300-400μ1. Polishing was done with a binding on a cobalt resin for 15min at 4°C, washed with buffer A and was eluted with 250mM of imidazole. The fractions were pooled and concentrated to 1-5 μM. Imidazole was removed by washing at concentration step.

### Nanobodies purification

Genes coding for C4 and E3 nanobodies (and E3 mutants) with cMyc tag and 6xHis-tag at the nanobody’s C-terminal were cloned into a pFUSE-derived vector (InvivoGen). The vector was used to transform Expi293F mammalian cells (ThermoFisher), and protein expression was carried out according to manufacturer’s recommendations using 100 ml final volume of cells suspension. Protein was then purified from the expression medium by affinity chromatography on a 1 mL HiTrap TALON crude column (Cytiva). After sample application, the column was first washed with 10 column volumes of PBS and 10 column volume of PBS supplemented with 10 mM Imidazole (pH=7.4) (Sigma), the protein was subsequently eluted with 5 column volumes of PBS supplemented with 0.5 M Imidazole (pH=7.4). Affinity-eluted nanobodies were finally polished on a HiLoad 16/600 Superdex 200pg Pre-packed column (Cytiva) using PBS buffer.

### Cryo-EM Sample Preparation

Typical grids were prepared with 1μM α7 pentamer supplemented with 10 or 50μM of nanobodies and 100μM nicotine when applicable. For preparation of cryo-EM grids, UltrAuFoil grids 300 mesh 1.2/1.3 or 0.6/1.0 from Quantifoil were glow discharged in a PELCO easiGlow for 25mA/15s. Three μl of protein were applied to grids before blotting for 6s at 100% humidity and 10°C. Grids were plunge-frozen with a Vitrobot Mark-IV (ThermoFisher). Frozen grids were stored in liquid nitrogen until use.

### Data collection, processing and model building

Datasets were collected on a ThermoFisher Glacios equipped with a Falcon 4i detector at Institut Pasteur, or a ThermoFisher Titan equipped with a Gatan energy filter bioquantum/K3 at Institut Pasteur or on a ThermoFisher IC-Krios equipped with a SelectrisX and a Falcon4i detector at EMBL-Heidelberg. All datasets were acquired in counting mode using EPU (ThermoFisher) or SerialEM (see Table supp 1 and Figures S4, S5, S9, S10, S11 for datasets specific details). The beam-induced motion was corrected by MotionCor2 and defocus values were estimated by CTFFIND-4.1. Laplacian-of-Gaussian algorithm from Relion4 or blobpicker from CryoSPARC3.3.2 were used to pick particles that were submitted to 2D classification. 2D classes showing landmark secondary structures of α7-nAChR were used to picked again particles. After two rounds of 2D classification, classes with clear secondary structures characteristic of α7 were selected as the initial pool of particles used for further processing. Unsymmetrized initial model was built in Relion4, symmetrized in C5 (or unsymmetrized in the case of C4partial) and used for Relion4 unsupervised 3D classification with a 60 Å initial low-pass filter. When indicated, an additional 3D classification without alignment was performed. The final pool of particles was used for 3D refinement in Relion4 with C5-symmetry imposed and relaxed in C1 to account for local heterogeneity, using a mask that includes signal we assigned to the transmembrane domain. FSC plots were calculated using the post-processing tool from Relion4.

Maps from the refinement runs were sharpened in Phenix1.20, except for the C4partial-Apo volume that was subjected to Phenix1.20 density modification using an initial model made with C4-Apo depleted in two nanobodies molecules. Low sigma value allows to visualize the transmembrane and intracellular regions, but they disappear at sigma values compatible with model building. On the first processed dataset (E3-Nic) molecular replacement tool from Phenix1.20 using the ECD of the Epi-bound nanodisc structure (PDB 7KOQ) was used to place α7. Phenix1.20 chain tracing was used to *de novo* build the five E3 molecules from their aminoacids sequence, except for the C-terminal Histag that is flexible and form a poorly resolved density on the apex of the structure. Model was modified in Coot and refined in Phenix1.20 using secondary structure and Ramachandran restraints. Using the glycosylation tool from Coot we were able to build the first sugars of the glycosylations on Asn23 and Asn67 and two sugars on Asn110 but densities suggest denser glycosylation trees. The two cysteine disulfide bonds in E3 were well resolved, as was the one in the α7 Cysloop. However, the two cysteines side chains from the canonic disulfide bond of the C-loop (Cys189/Cys190), although well resolved, do not appeared bond in the density. This loop is exposed to the solvent and the disulfide bond is known to be sensitive to radiation damage (Rhaman et al, Noviello et al, Hattne et al 2018), we thus omitted the bond in the structure. For the remaining datasets, the resulting E3-Nic structure was used for molecular replacement, sequence of the nanobodies was further ajusted for C4 and structures were refined using Coot and Phenix and validated using the MolProbity Phenix validation tool.

Structure analysis was performed using Pymol, ChimeraX and PISA from the CCP4 suite.

### Real-time surface plasmon resonance assays

SPR assays were performed on a Biacore T200 instrument (Cytiva) equilibrated in buffer A pH8 at 25°C.

Surface preparation: Carboxy-methylated dextran CM5 sensorchips (Cytiva) were functionalized by immobilizing covalently the anti-Rho tag monoclonal antibody 1D4 (50μg/ml at pH4) through amide bonds at a density of 13000-15000 resonance units (RU, lRU≈lpg/mm2). α7-nanodiscs (10μg/ml) were then captured on the 1D4 surfaces at a density of 1300-1500RU.

Binding assays: the association and dissociation kinetic properties of nanobodies/α7 complexes were determined by injecting nanobodies in single cycle kinetics mode (five increasing concentration injections of 600s each at 30μl/min on both α7/Rho1D4 and reference Rho1D4-only surfaces. This was followed by a final dissociation phase of 1800s. Three-fold dilution series were used for each nanobody (100-1.23 nM and 20-0.25nM for E3 and its mutants; 6.66-0.08nM for C4). The specific SPR signals were analysed using the Biacore T200 evaluation software (Cytiva) yielding association (k_on_) and dissociation (k_off_)rates, and equilibrium dissociation constants (K_d_) for each nanobody/α7 complex.

### Immunofluorescence

HEK293 cells were cultured on poly-D-lysine (Millipore) coated glass coverslips, according to manufacturer’s recommendations. The hα7-nAChR-IRES-eGFP transfer plasmid or its mutated version (α3MIR or S25A) was co-transfected with NACHO, Ric-3 and SAT1 plasmids (1:1:1:1 ratio) using and the JetPrime transfection reagent (Polyplus), according to manufacturer instructions.

48h-36h after transfection, cells were fixed with 4% PFA and permeabilized for 2 min with a solution of ethanol/methanol (1:1). Non-specific binding was blocked with 10% BSA in PBS for 5 min at room temperature. The nanobodies-Fc constructs were diluted to 5μg/ml in 10% BSA in PBS and incubated with the coverslips for 2h at room temperature.

Proper expression of α7-nAChRs was verified by a mouse anti-Rho1D4. Anti-human IgG and anti-mouse IgG coupled to Alexa647 (ThermoFisher) were diluted in PBS-BSA.

Coverslips were mounted on slides after Prolong-DAPI staining (Invitrogen) and visualized using epi-fluorescence at constant exposure times. All experiments were reproduced ≥4 times.

### Two-electrode voltage-clamp electrophysiology

*Xenopus laevis* oocytes were obtained from EcoCyte Bioscience, Germany and from Tefor Paris-Saclay UAR2010 and maintained in modified Barth’s medium (87.34 mM NaCl, 1 mM KCl, 0.66mM NaNO_3_, 0.75 mM CaCl_2_, 0.82mM MgSO_4_, 2.4 mM NaHCO_3_, 10 mM HEPES pH 7.6). Defolliculated oocytes were submitted to intranuclear injection of ~2–6 ng of hα7-nAChR-IRES-eGFP transfer plasmid and kept at 18 °C for 2–3 days before recording.

Recordings were performed with a Digidata 1550A digitizer (Molecular Devices), an Axon Instruments GeneClamp 500 amplifier (Molecular Devices), an automated voltage-controlled perfusion system which controls an 8-port and a 12-port electric rotary valves (Bio-Chem Fluidics) both connected to a 2-way 4-port electric rotary valve (Bio-Chem Fluidics) and the pClamp 10.6 software (Molecular Devices).

Oocytes were perfused with Ringer’s buffer (100 mM NaCl, 2.5 mM KCl, 10 mM HEPES, 2 mM CaCl_2_, 1 mM MgCl_2_, pH 7.3). Nanobodies solutions were applied after dilution in Ringer’s buffer and all currents were measured at - 60mV. For all the experiments oocytes were perfused with Ringer’s buffer for 30s, then 5s with 30μM ACh (or 0,3μM for L9’T) followed by 2- or 3-minutes wash and another 5s ACh application. The nanobody solution was then perfused for 30s (or 10s for L9’T) followed by 5s of 30μM ACh and 2- or 3-minutes wash and another 5s ACh application.

Recordings were analyzed using ClampFit and GraphPad Prism. Measurements were performed at the peak of the response. For the statistical comparisons using GraphPad Prism, we performed Student’s *t test*.

## Acknowledgements

This work was supported by an ERC (Grant no. 788974, Dynacotine) to PJC and MSP, an Institut Pasteur Axe3 Seed grant to MSP, iNEXT-Discovery funding (PID: 22438) to MSP, the Foundation de la Recherche Médicale (Equipe FRM DEQ20140329497) to PJC, the Agence Nationale de la Recherche (Grant ANR-21-CE37-0026, Nicoptotouch) to PJC and GA, and the Institut National du cancer INCA to GBD. The authors thank the Nanoimaging Core Facility (Institut Pasteur) and Simon Fromm (EMBL-HD) for their precious help in data collection, and H. Nury, A. Menny and N. Wolff for discussions and critical reading of the manuscript.

## Authors contributions

MSP and PJC conceived and supervised the study. NB and SP prepared lentiviruses with UM supervision. PL and GA produced nanobodies. MSP and NB setup and performed protein production, cryoEM samples preparation and data collection. MG, GPA and FB helped in early biochemistry, electron microscopy and data processing respectively. MSP processed the cryoEM data and solved the structures. NB and PE performed SPR experiments. GDB performed electrophysiology experiments. All authors helped in the results analysis and interpretation. MSP and PJC wrote the manuscript with contributions from all authors.

## Figures supp

**Supplementary figure 1:**
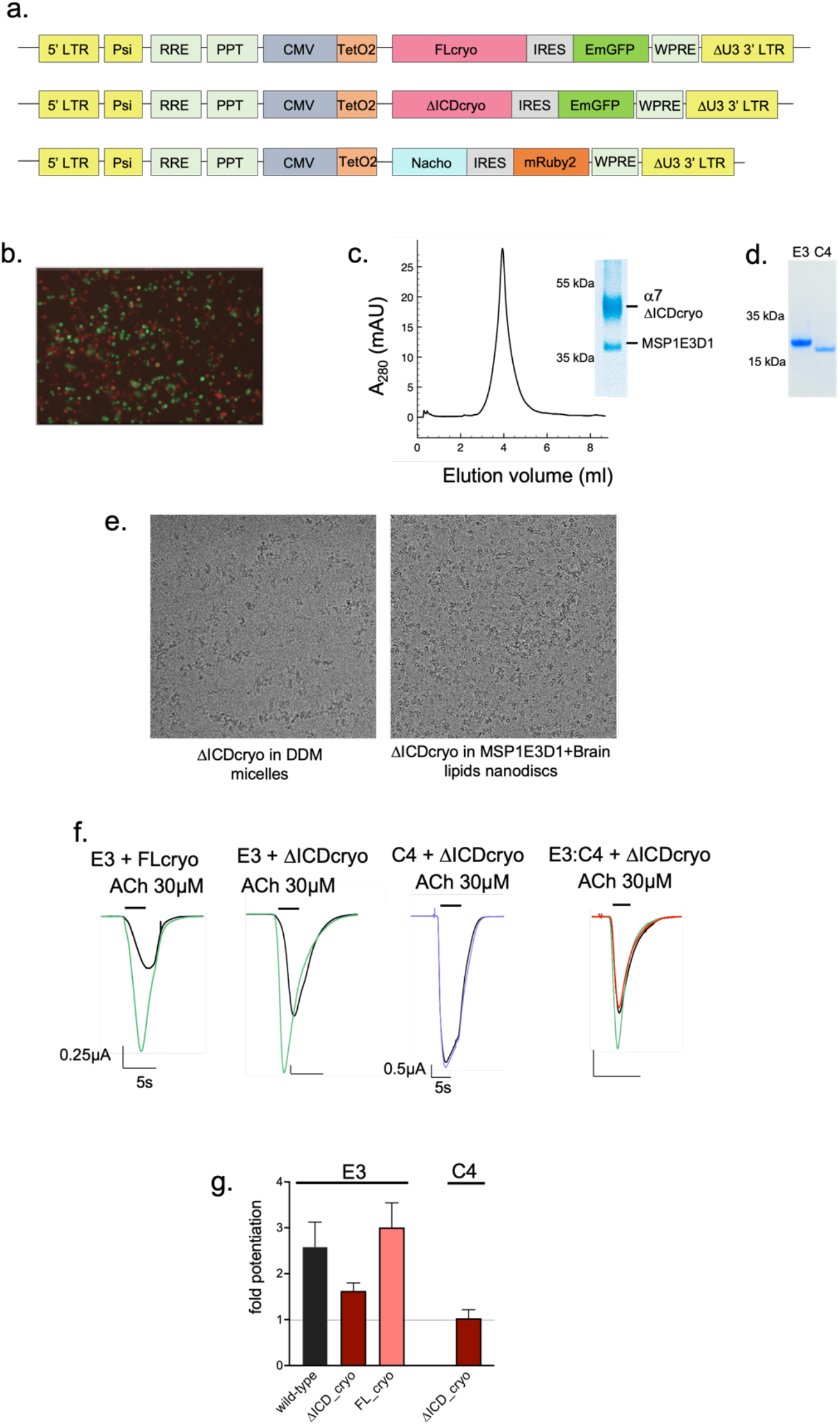
Production of the α7 nAChR and functional characterization of E3 WT and mutants. a. Lentiviral transfer regions used in the study. LTR: Long Terminal Repeat sequence, ΔU3: Deletion in 3’LTR rendering the virus “self-inactivating” (SIN) after integration, Psi : Packaging signal sequence; RRE: Rev Response Element cPPT: Central Polypurine Tract; CMV: Cytomegalovirus Promoter; TetO2 : Tetracycline-controlled transcriptional activation sequence; IRES : Internal ribosome entry site of the encephalomyocarditis virus; WPRE: Woodchuck Hepatitis Virus Post-transcriptional Regulatory Element b. After transduction and induction, most cells exhibit orange and green fluorescence indicating efficient cell infection. c. Size-exclusion profile of α7ΔICDcryo reconstituted in nanodiscs and SDS PAGE of the concentrated sample before plunge freezing. d. SDS-PAGE of E3 and C4 after purification. e. Representative micrographs of α7ΔICDcryo in detergent micelles or reconstituted in nanodiscs, showing the presence of aggregates in the former. f. Representative TEVC recordings showing the potentiation of 30μM ACh-elicited currents upon a 30s pre-application of 1μM of E3 (green) or C4 (purple) or by 0.25μM E3 and 0.5μM C4 (red) with the indicated α7 constructs. g. Fold potentiation of 30μM ACh currents determined with a 30sec pre-application of 1μM nanobody for the indicated constructs. Data are mean ±sd for n≥4.

**Supplementary figure 2:**
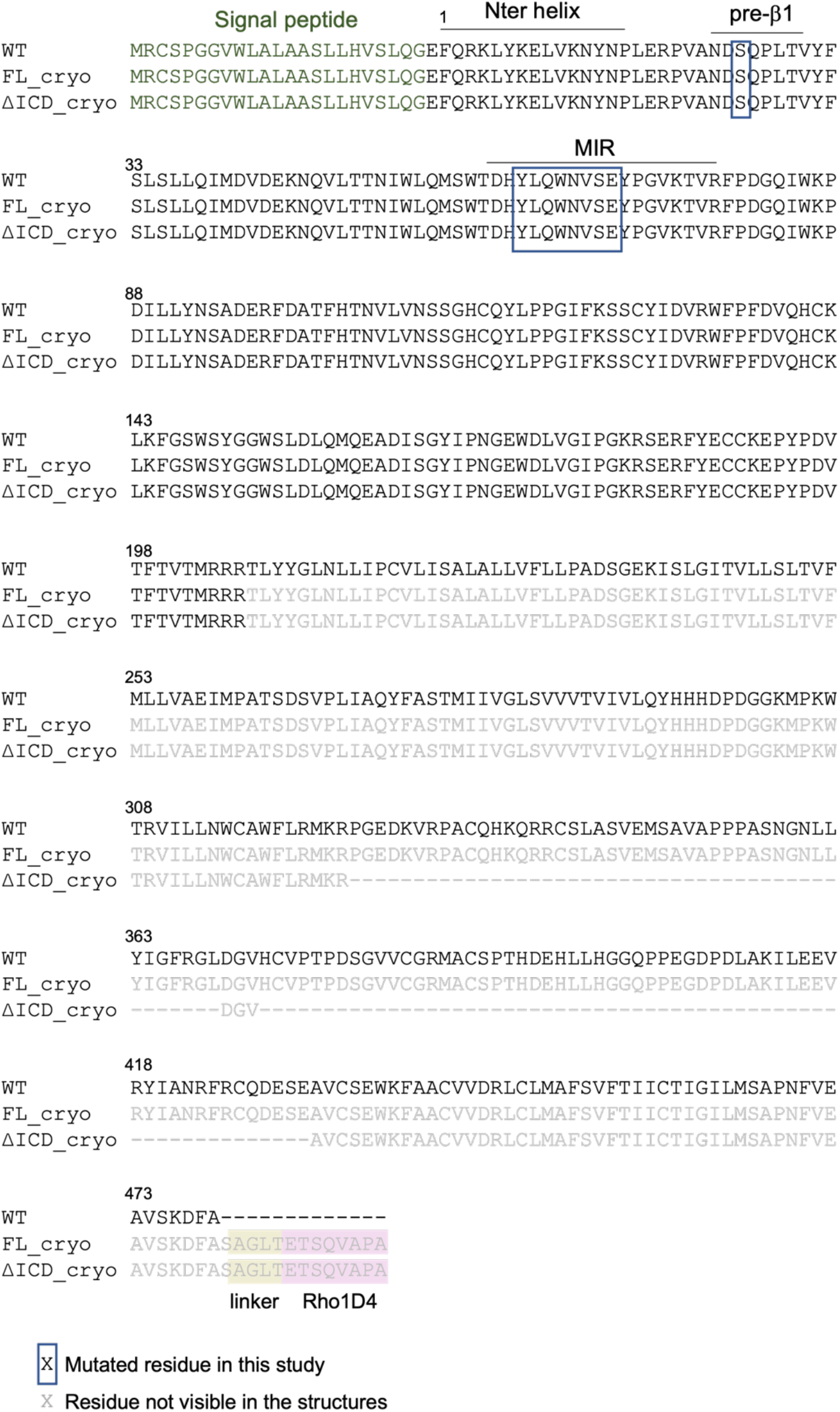
Sequence alignment of human α7-nAChRs and the constructs used in this study. Signal peptide regions are denoted in green. A box locates the residues mutated in this study. Grey residues form the transmembrane and intracellular regions not resolved in the structures.

**Supplementary figure 3:**
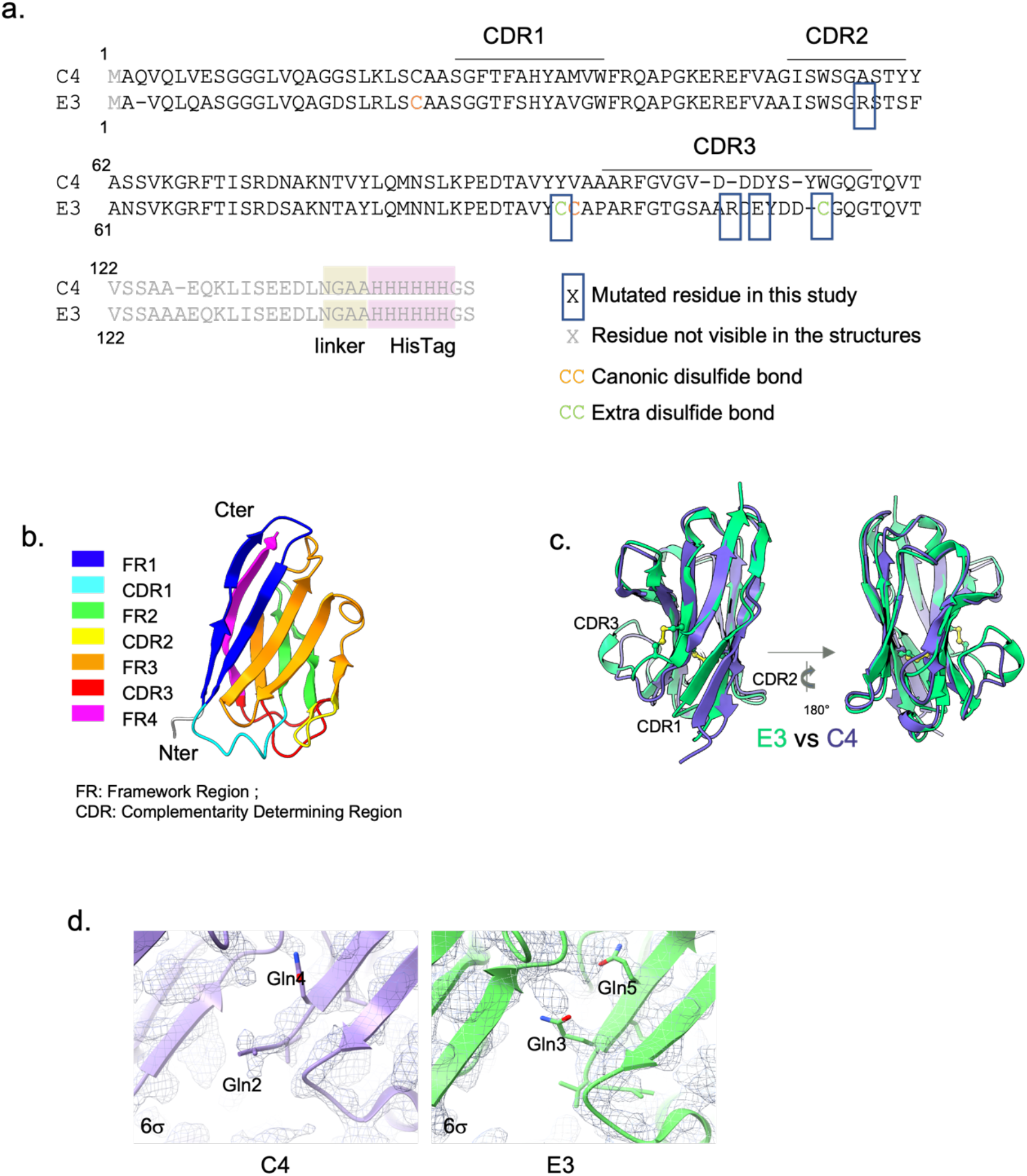
Sequence and structure of nanobodies E3 and C4. a. Sequence alignment of E3 and C4. A box locates the residues mutated in this study. The cysteines involved in the canonical and extra disulfide bonds are denoted in orange and green respectively. Grey residues are not resolved in the structures. Note the missing Gln at the N-terminus of E3 that shifts the numbering of the whole nanobody compared to C4. b. Topology of the nanobodies exemplified with the structure of C4. Elements are rainbow-colored from the Nter (blue) to Cter (pink). c. Tertiary structure comparison of C4 (purple) and E3 (green). The two disulfide bonds of E3 are shown in sticks representation. d. The N-terminus of C4 extends towards the solvent while the N-terminus of E3 folds back inside the β-sandwich. Close-view of the Nter of the two nanobodies as seen in the C4-Apo and E3-Apo datasets and their corresponding electron densities contoured at 6σ. The two aligned glutamine residues are shown in sticks.

**Supplementary figure 4:**
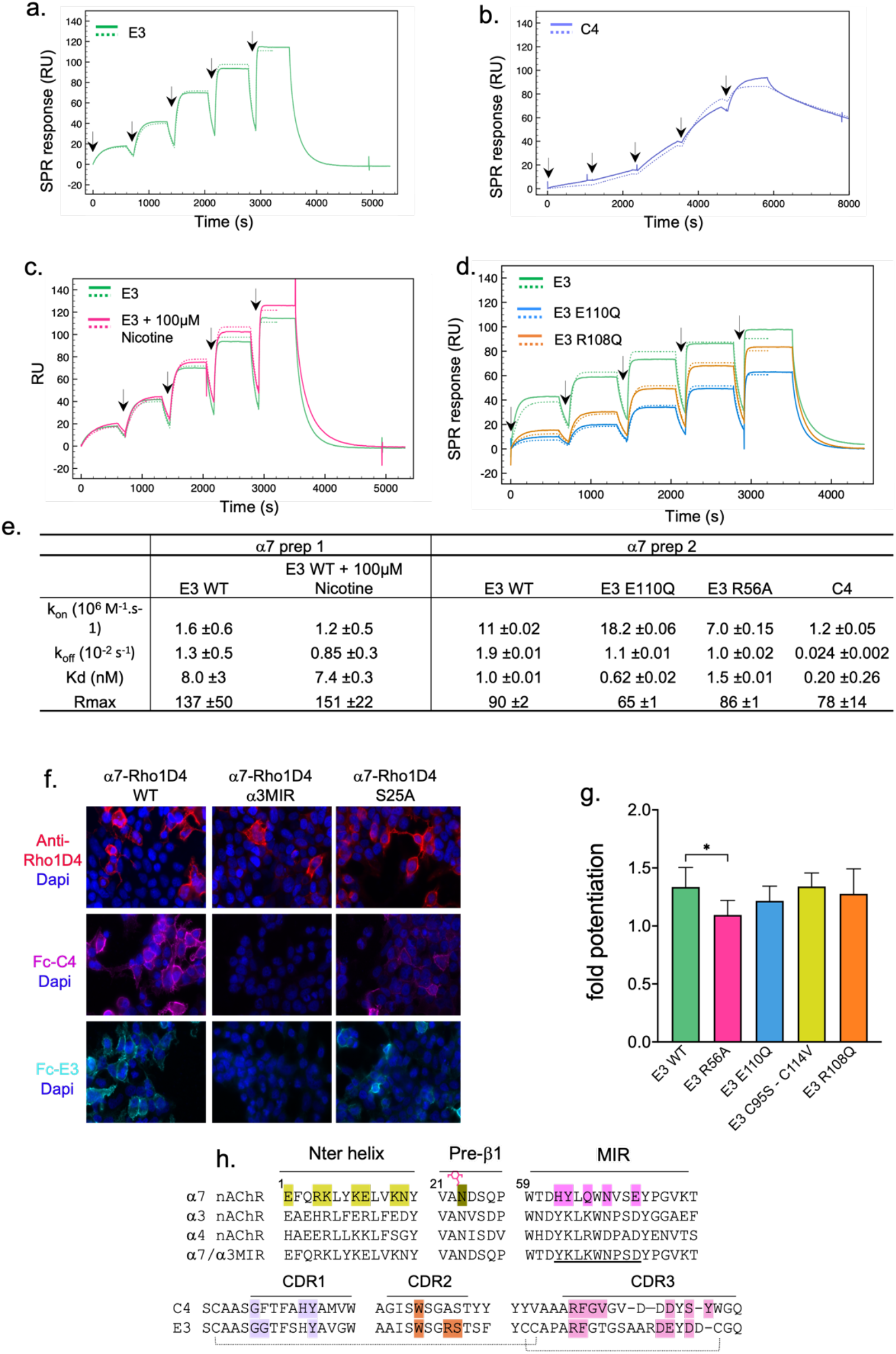
characterization of the α7 nAChR, C4, E3 and their mutants by SPR, immunofluorescence and electrophysiology. a. Real-time SPR monitoring of injections of increasing concentrations of E3 (black arrows) on purified α7ΔICDcryo reconstituted in nanodiscs. The dashed line shows the fitting curve used to determine the kinetic constants. b. Same as in a. with C4. c. Same as in a. with E3 (green) and E3 with 100μM nicotine in the running buffer (pink). d. Same as in a with E3 (green), E3 E110Q (blue) and E3 R108Q (orange). e. Kinetic and affinity constants determined by single cycle kinetics surface plasma resonance. E3 WT was first assayed with or without 100μM nicotine on a single α7ΔICDcryo nanodiscs preparation. Another preparation was used to perform the assay with E3 WT and its mutants and with C4. Data are mean ±sd of 3 independent channels on the sensorchip. f. Immunofluorescence of C4 and E3 fused to Fc on HEK cells expressing α7 WT, S25A or the α7/α3MIR chimera. Top: α7 receptors are labelled using the Cter Rho1D4 tag (red), nuclei using Dapi (blue). Middle: Fc-C4 labelling is shown in purple, nuclei are labelled using Dapi (blue). Bottom: Fc-E3 labelling is shown in green, nuclei are labelled using Dapi (blue). g. Fold potentiation calculated by TEVC on n≥4 cells by E3-WT and its mutants on α7-WT. Values were submitted to a paired t-test that is denoted with * p≤0.05 when a significant difference was found. h. Sequence alignments of the loops involved in binding. Top: alignment of the Nter helix, pre-β1 and MIR of α7, α3, α4 and the chimeric α7/α3MIR construct (with the chimeric sequence underlined). Key residues involved in binding are highlighted and the glycan on Asn23 depicted as a pink sugar. Sequence numbering is the one of α7. Bottom: alignments of the three CDR regions of C4 and E3. Key residues involved in binding are highlighted. Disulfide bonds of E3 are depicted by links between the two cysteines. CDR3s alignment is based on their local structure.

**Supplementary figure 5:**
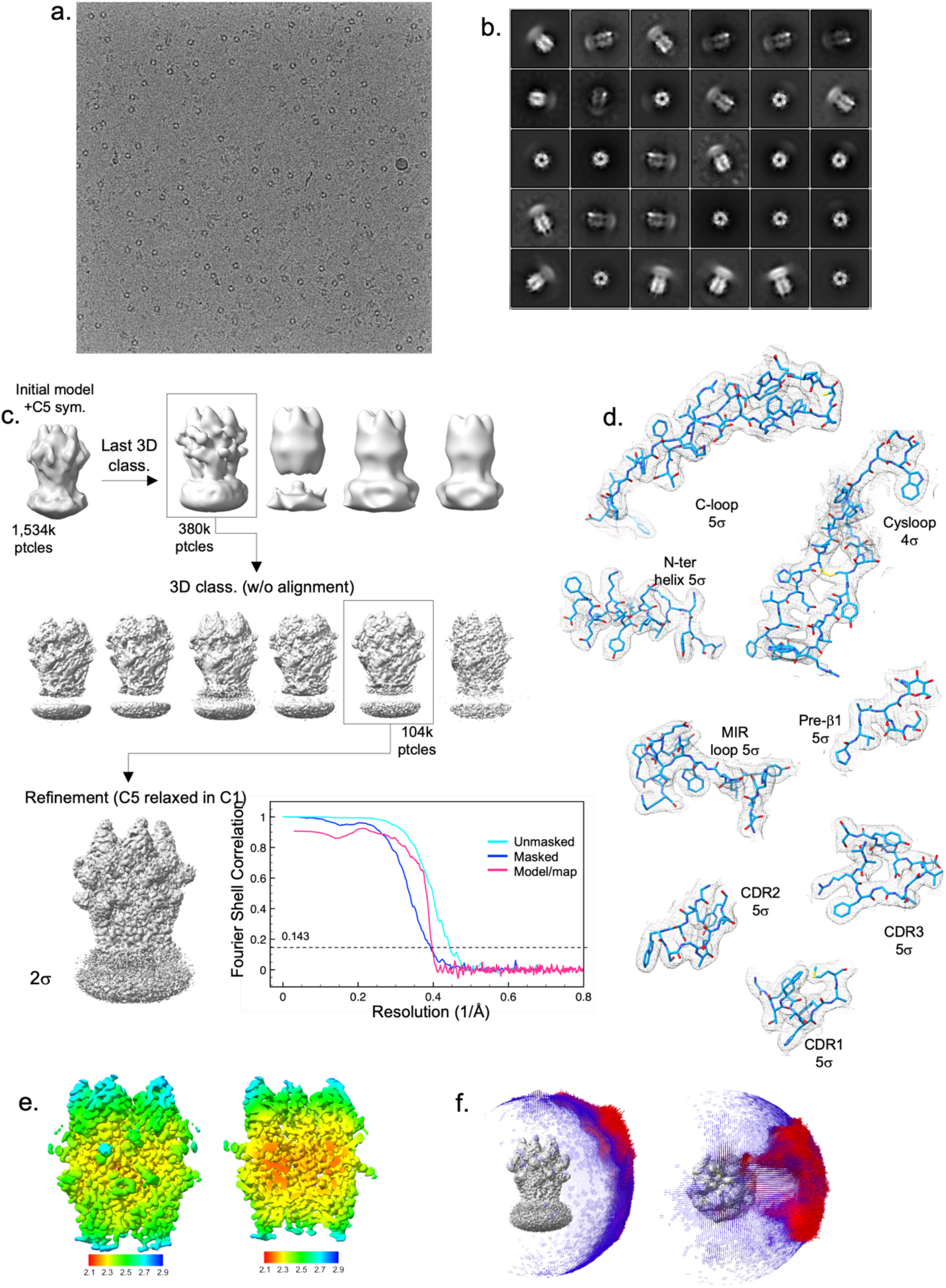
Electron microscopy and 3D reconstruction of the C4-Apo data set. a. Representative micrograph of the C4-Apo dataset. b. Selected 2D class averages c. Flowchart of the 3D volume processing. A C5-symmetry was applied on the initial model for 3D classification with particles alignment. The best class particles were submitted to another 3D classification round without alignment and high-resolution reaching particles further refined with a C5 symmetry relaxed in C1 (Relion 3D refine). Resulting volume is shown at 2σ contouring with the FSC curves on its right. d. Representative densities and model building for the loop C, cys-loop, Nter helix, pre-β 1, MIR of α7 and the three CDRs of C4 with the indicated contouring level e. Local resolution map calculated with Relion. The volume is shown at high contouring from the side and sliced in the vestibule. f. Angular distribution of the particles seen on the side and top views of the unsharpened map

**Supplementary figure 6:**
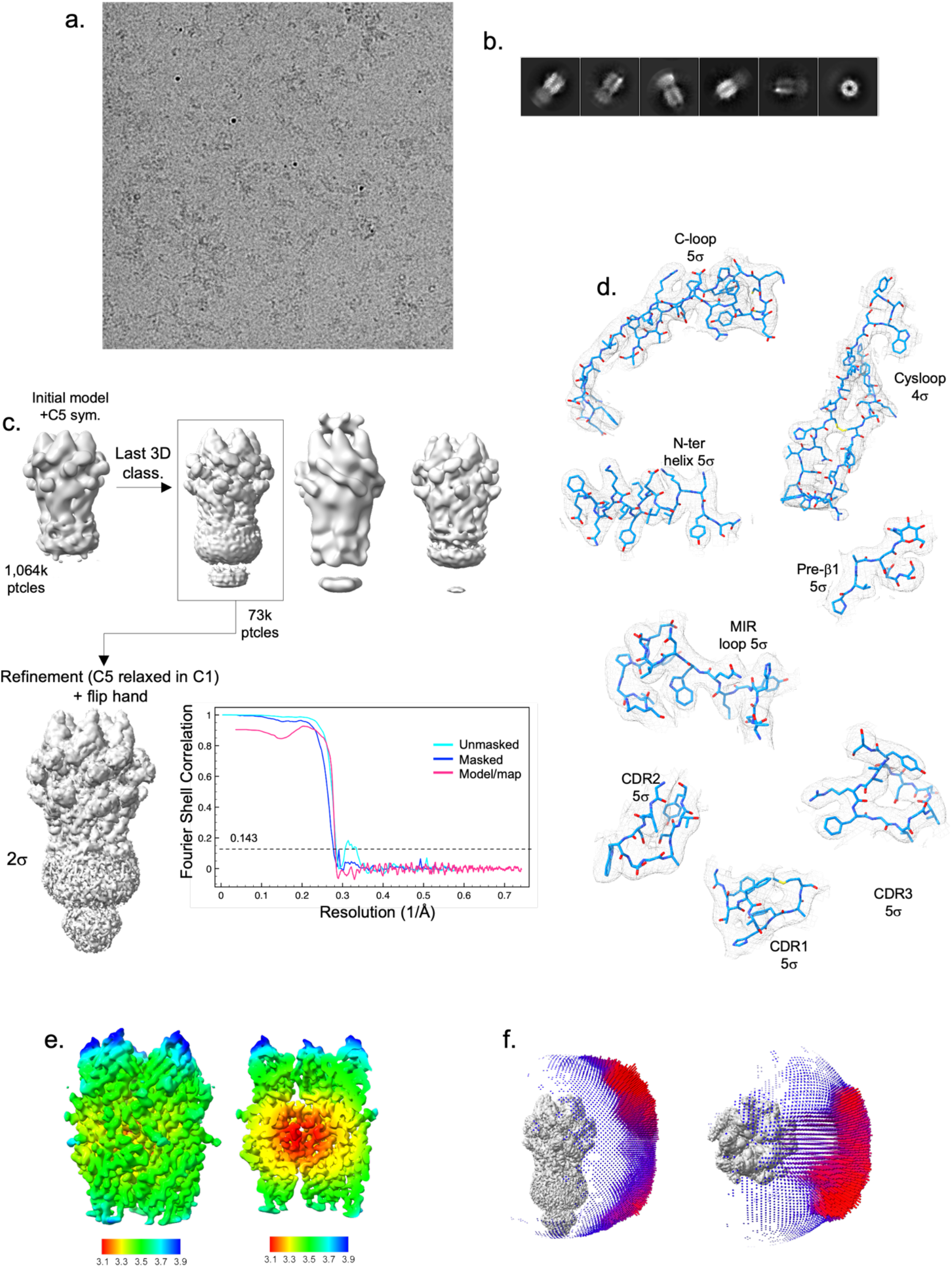
Electron microscopy and 3D reconstruction of the C4-Nic data set. a. Representative micrograph of the C4-Nic dataset. b. Selected 2D class averages c. Flowchart of the 3D volume processing. A C5-symmetry was applied on the initial model for 3D classification with particles alignment. The best class particles were further refined with a C5 symmetry relaxed in C1 (Relion 3D refine). Resulting volume is shown at 2σ contouring with the FSC curves on its right. d. Representative densities and model building for the loop C, cys-loop, Nter helix, pre-β1, MIR loop of α7 and the three CDR of C4 with the indicated contouring level e. Local resolution map calculated with Relion. The volume is shown at high contouring from the side and sliced in the vestibule. f. Angular distribution of the particles seen on the side and top views of the unsharpened map

**Supplementary figure 7:**
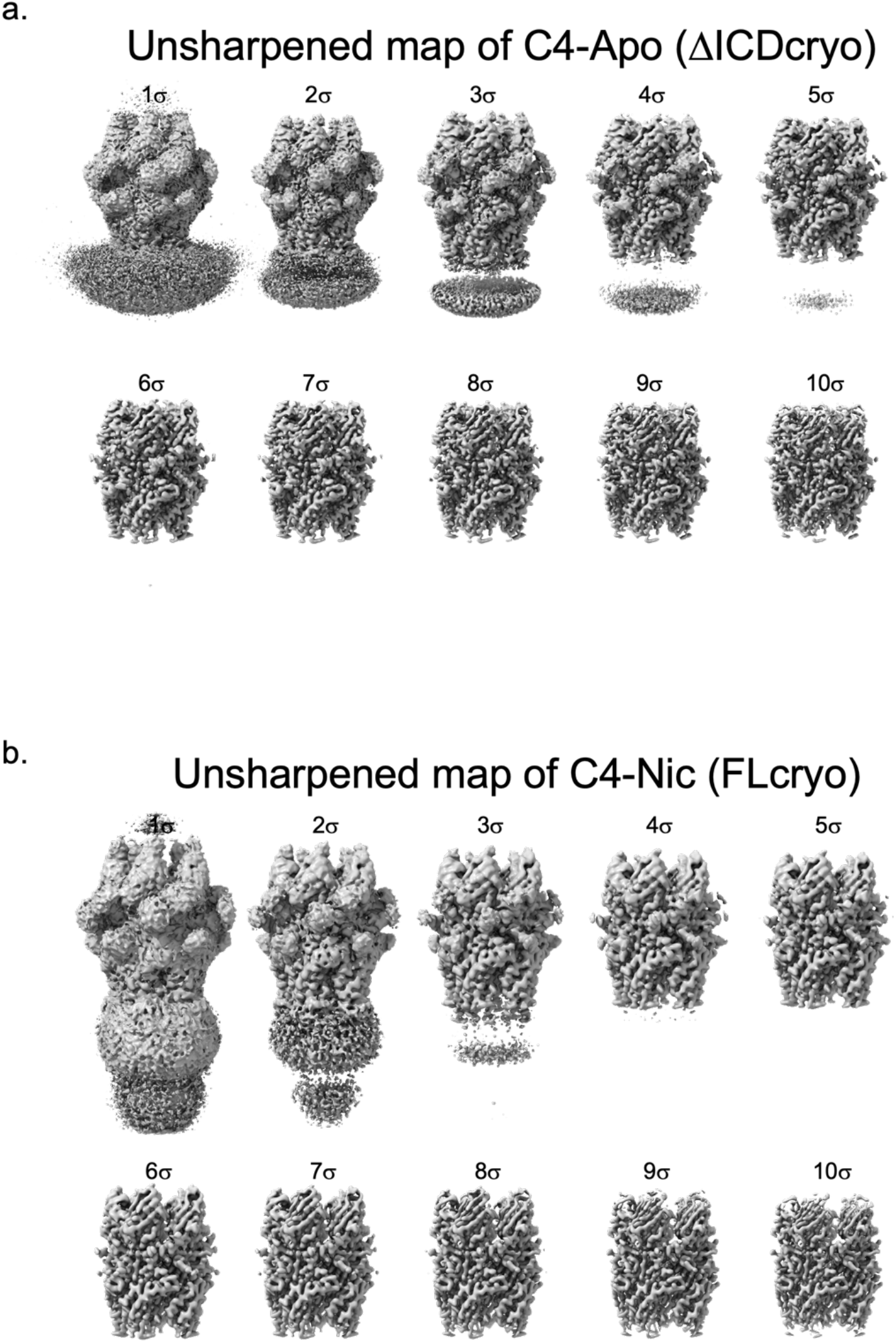
Unsharpened cryo-EM maps of C4-Apo and C4-Nic. a. Various contouring levels applied to the C4-Apo density showing the flexibility of the transmembrane/nanodisc region that completely disappears at 6σ. b. Various contouring levels applied to the C4-Nic density showing the flexibility of the transmembrane/nanodisc and intracellular regions that completely disappear at 4σ.

**Supplementary figure 8:**
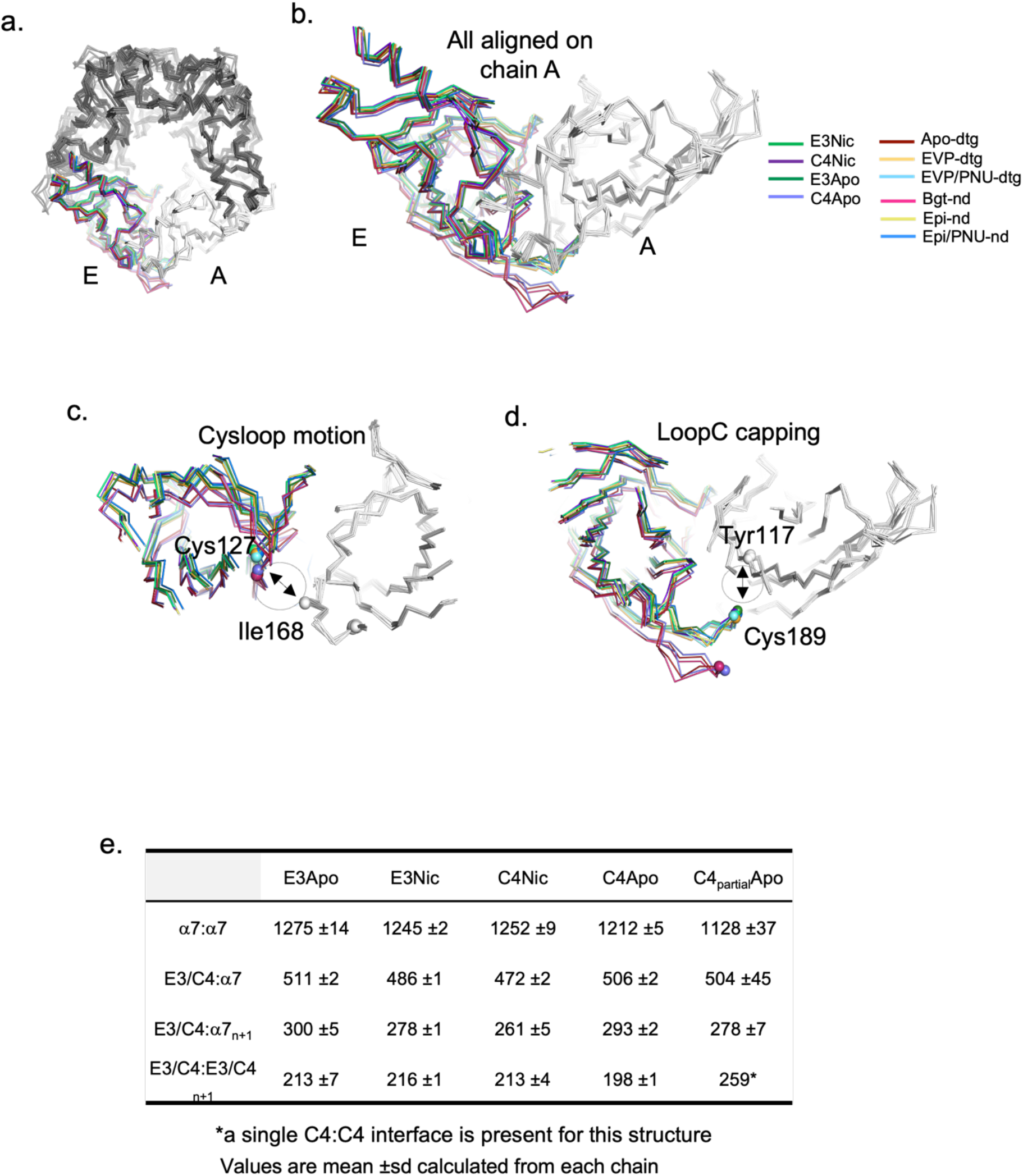
Analysis of the conformation of the α7 ECD in E3 and C4 co-structures. a. α7 pentamers from known structures are aligned at the level of their chain A and represented as ribbons in top view. b. A dimer A-E of the aligned structures in a. is shown. Chains A are seen in grey and chains E according to the color-coding on the right. c. Same as in b. but clipped to reveal the cys-loop and its motion towards the chain A. The “cys-loop motion” distance measurement between the Cα of Cys127 and Ile 168 is figured with an arrow. d. Same as in b. but clipped to reveal the Loop C and its motion towards the chain A. The “Loop C capping” distance measurement between the Cα of Cys189 and Tyr117 is figured with an arrow. e. Interaction surface areas calculated using PISA (CCP4 suite) in Å ^2^.

**Supplementary figure 9:**
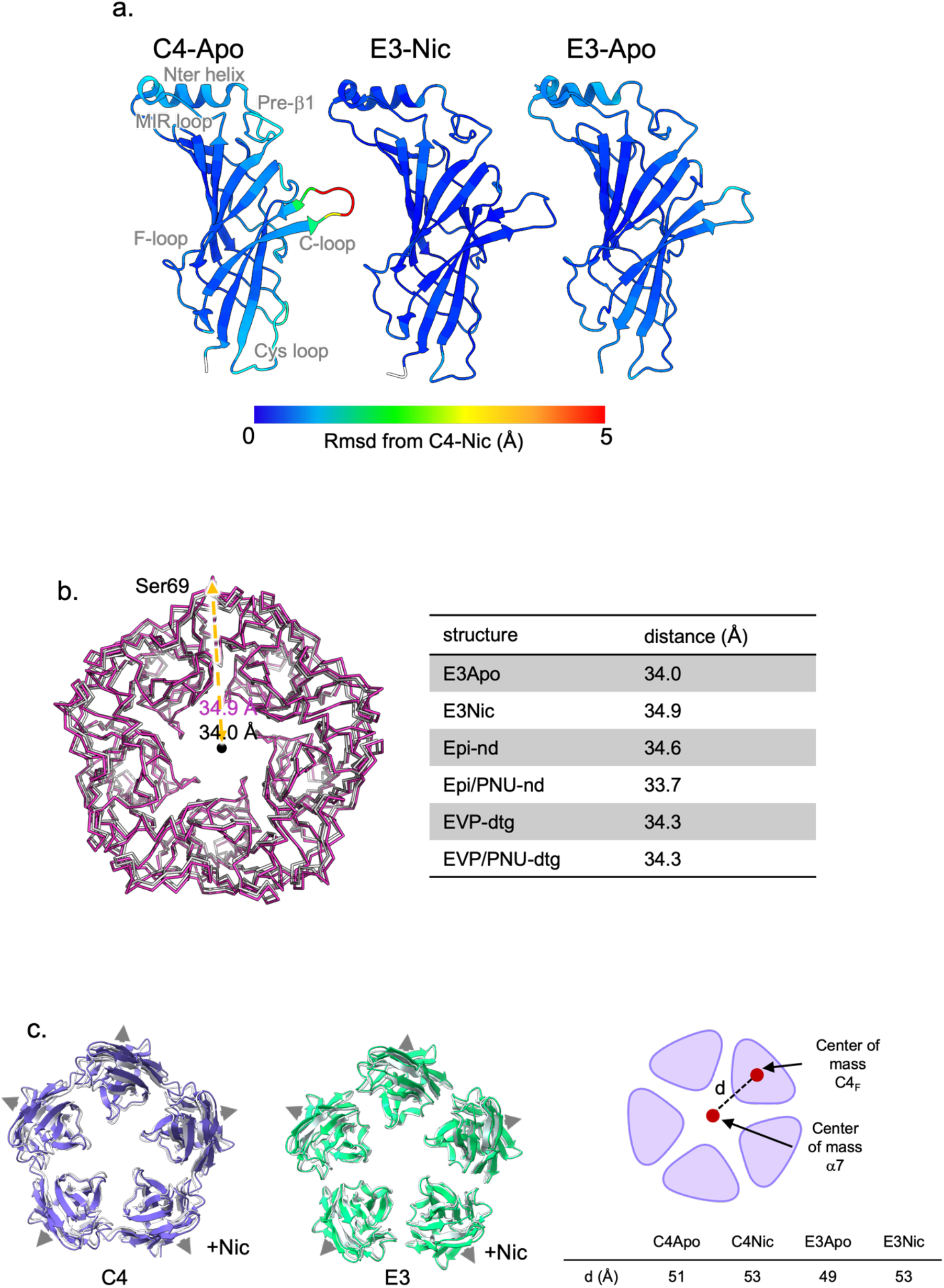
Tertiary and quaternary reorganization of nanobody-bound structures in the presence of nicotine. a. Tertiary variations of α7 between structures. Cα rmsd were calculated in ChimeraX using the C4-Nic structure as reference upon superimposition of monomers. ECD monomer of C4-Apo, E3-Nic and E3-Apo are shown in cartoons were Cα are colored according to the rmsd value, with the same range for all and shown in the color key. b. ECD expansion between E3-Apo and E3-Nic. Distances are measured between the Cα of Ser69 from chain A and the center of mass of the Cα of the five Ser69 which figure the middle of the upper part of the ECD. The distance is shown on a top view of the E3-Apo (grey) and E3-Nic (pink) aligned and represented in ribbons. The table shows the values of this distance on several structures. Note that the expansion is absent between the two detergent structures and is similar for the two pairs of nanodisc structures. c. Nanobodies expansion upon nicotine binding. With both nanobodies, we observed a small outward motion of the nanobodies molecules upon nicotine binding, as represented from the top with Apo structures in white and nicotine-bound structures in purple and green respectively. We found this motion to be around 2-3 Å by measuring the distance between the center of mass of one nanobody and the center of mass of α7 (summarized in the table).

**Supplementary figure 10:**
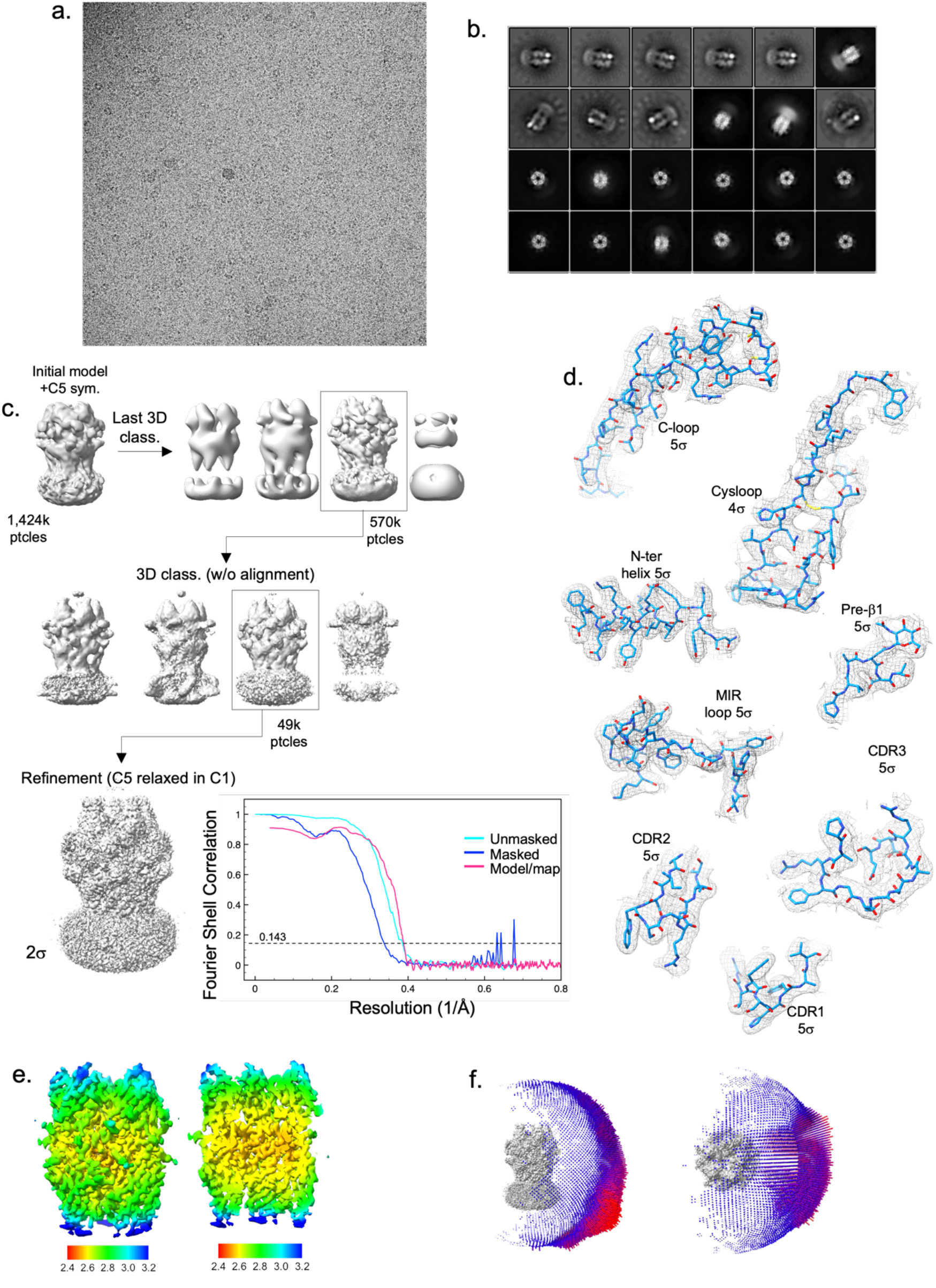
Electron microscopy and 3D reconstruction of the E3-Apo data set. a. Representative micrograph of the E3-Apo dataset. b. Selected 2D class averages c. Flowchart of the 3D volume processing. A C5-symmetry was applied on the initial model for 3D classification with particles alignment. The best class particles were submitted to another 3D classification round without alignment and high-resolution reaching particles further refined with a C5 symmetry relaxed in C1 (Relion 3D refine). Resulting volume is shown at 2σ contouring with the FSC curves on its right. d. Representative densities and model building for the loop C, cys-loop, Nter helix, pre-β 1, MIR of α7 and the three CDRs of E3 with the indicated contouring level. e. Local resolution map calculated with Relion. The volume is shown at high contouring from the side and sliced in the vestibule. f. Angular distribution of the particles seen on the side and top views of the unsharpened map.

**Supplementary figure 11:**
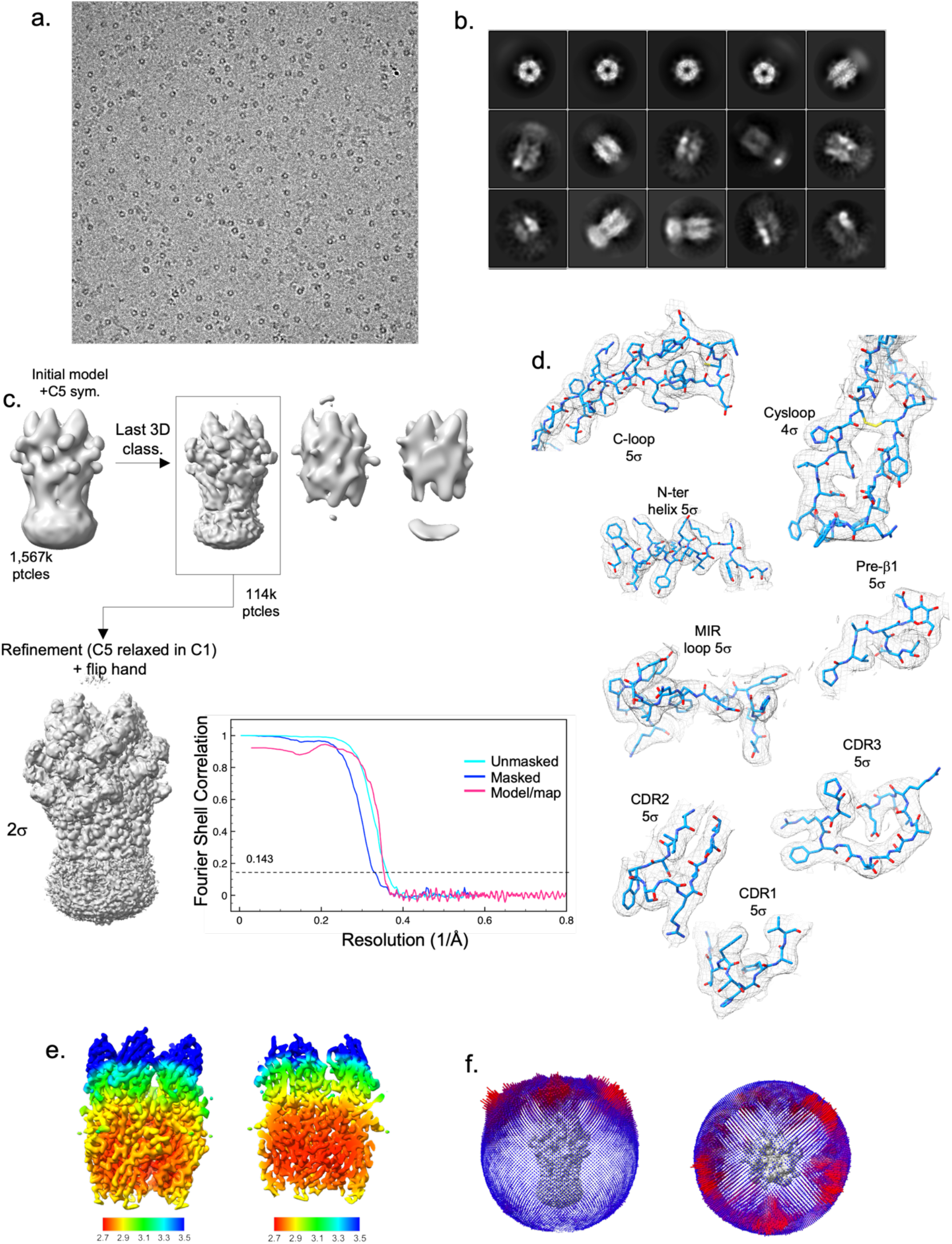
Electron microscopy and 3D reconstruction of the E3-Nic data set. a. Representative micrograph of the E3-Nic dataset. b. Selected 2D class averages c. Flowchart of the 3D volume processing. A C5-symmetry was applied on the initial model for 3D classification with particles alignment. The best class particles were further refined with a C5 symmetry relaxed in C1 (Relion 3D refine). Resulting volume is shown at 2σ contouring with the FSC curves on its right. d. Representative densities and model building for the loop C, cys-loop, Nter helix, pre-β 1, MIR of α7 and the three CDRs of E3 with the indicated contouring level. e. Local resolution map calculated with Relion. The volume is shown at high contouring from the side and sliced in the vestibule. f. Angular distribution of the particles seen on the side and top views of the unsharpened map.

**Supplementary figure 12:**
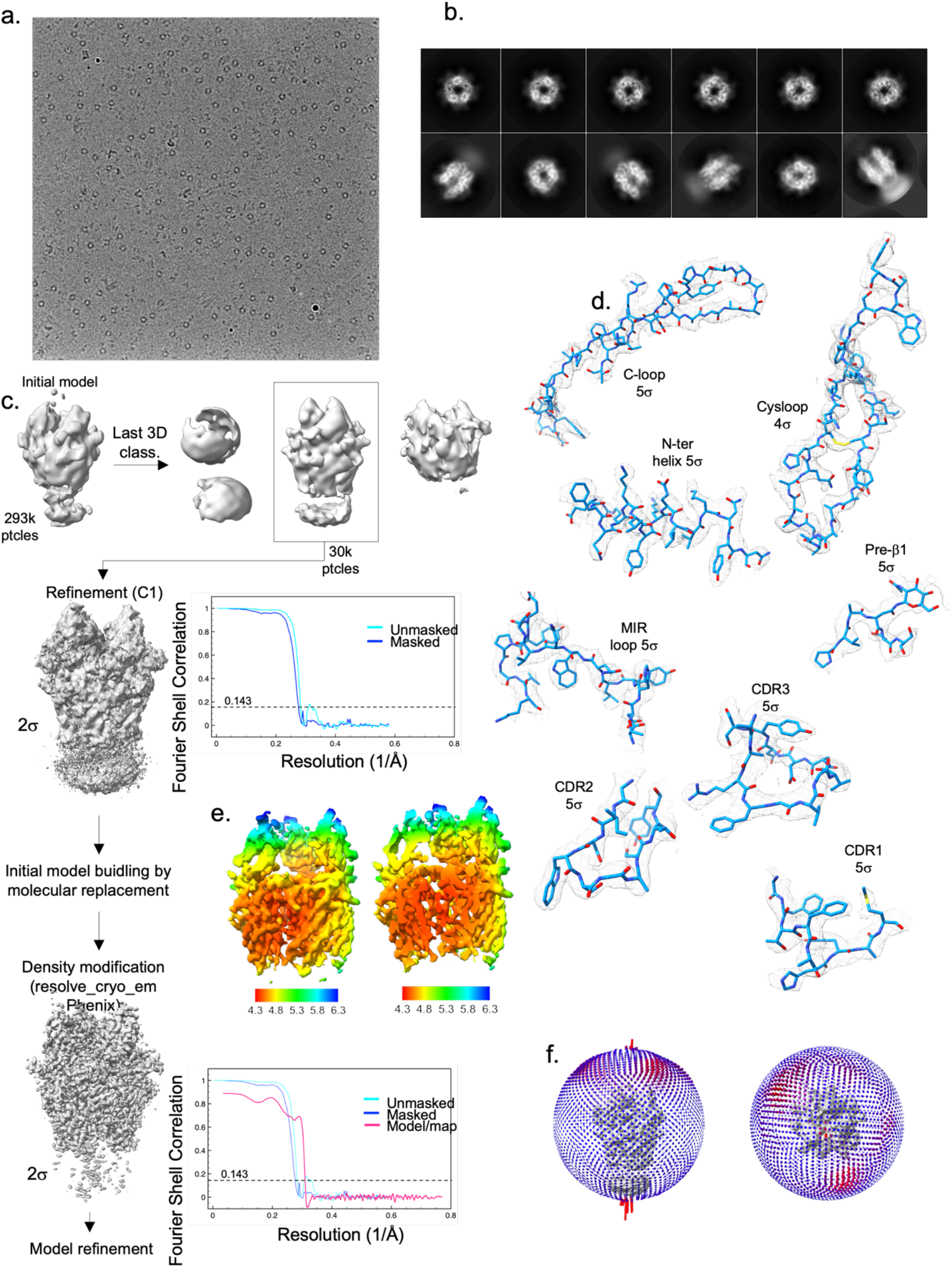
Electron microscopy and 3D reconstruction of the C4partial-Apo data set. a. Representative micrograph of the C4partial-Apo dataset b. Selected 2D class averages c. Flowchart of the 3D volume processing. No symmetry was applied on the initial model for 3D classification with particles alignment. The best class particles were further refined (Relion 3D refine). Resulting volume is shown at 2σ contouring with the FSC curves on its right. After a first model building, the density and the model were subjected to density modification in Phenix, resulting in a volume allowing the refinement of a protein model at 3.4 resolution shown at 2σ contouring with the FSC curves on its right d. Representative densities and model building for the loop C, cys-loop, Nter helix, pre-β 1, MIR of α7 and the three CDRs of C4 with the indicated contouring level. e. Local resolution map calculated with Relion. The volume is shown at high contouring from the side and sliced in the vestibule. f. Angular distribution of the particles seen on the side and top views of the unsharpened map.

**Supplementary figure 13:**
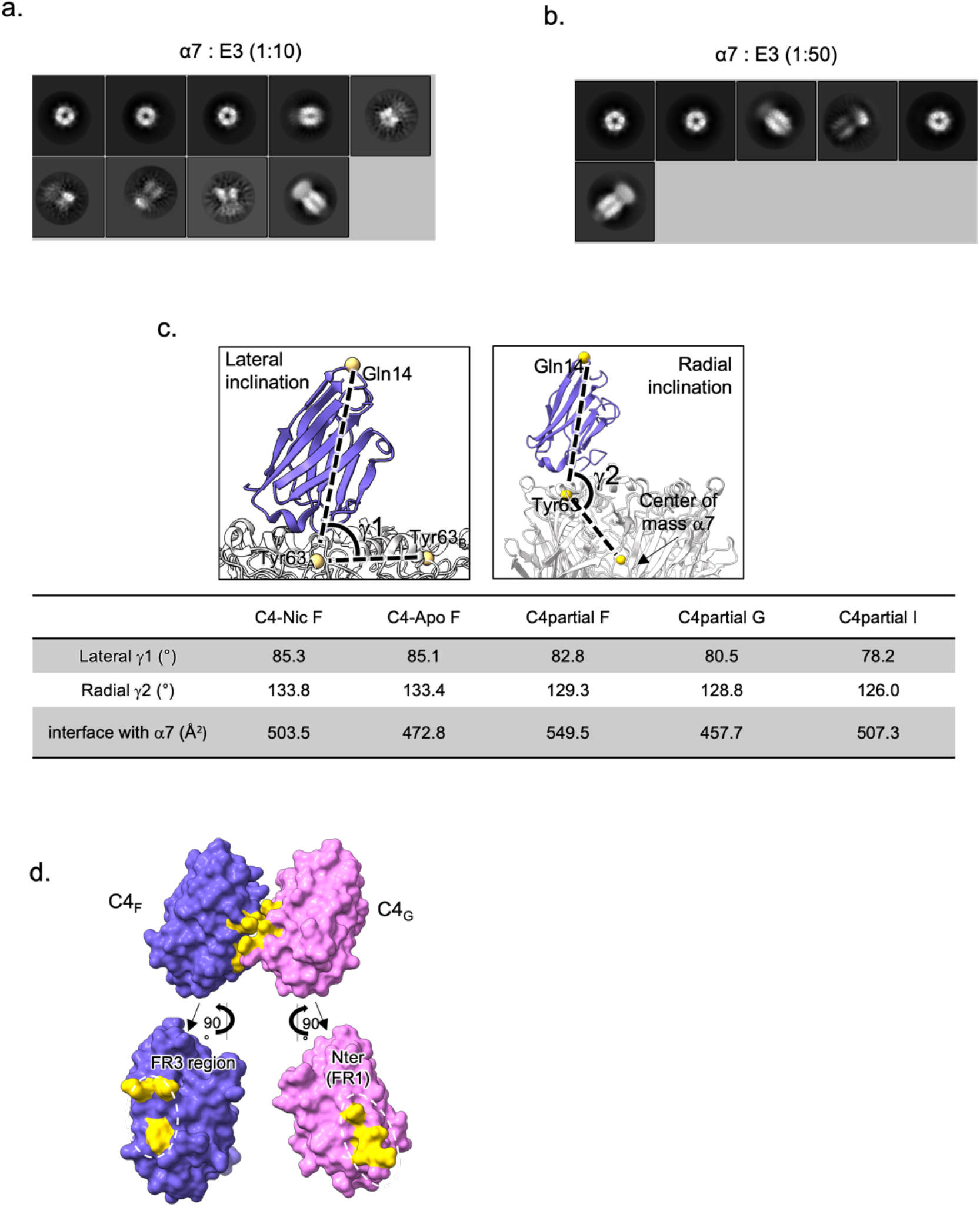
Analysis of the C4partial-Apo dataset. a. Preliminary work was done with lower concentration of E3 in the grids (10μM) and early 2D classes showed heterogeneity with most classes showing 4 molecules bound to α7. b. Grids prepared the same day with 50μM E3 showed a vast majority of particles with 5 E3 molecules bound. c. Orientation of the C4 molecules in the C4partial-Apo structure. The lateral inclination angle is defined between the Cα of Gln14 in C4, of Tyr63 in the α7 chain below and of Tyr63 of the complementary α7 subunit. The radial inclination angle is defined between the Cα of Gln14 on C4, of Tyr63 on the α7 subunit below and the center of mass of the α7 pentamer. d. Surface representation of the two adjacent C4 (F in purple, G in pink) in the C4partial-Apo dataset and their exploded view below. Residues at the interface are colored in yellow.

**Supplementary Table 1.** Cryo-EM data collection, refinement and validation statistics.

|  | C4-Apo<br>(EMDB-)<br>(PDB ) | C4-Nic<br>(EMDB-(PDB | E3-Apo<br>(EMDB)<br>(PDB ) | E3-Nic<br>(EMDB)<br>(PDB ) | C4partial-Apo<br>(EMDB)<br>(PDB ) |
| --- | --- | --- | --- | --- | --- |
| **Data collection and processing** |  |  |  |  |  |
| Microscope | IC-Krios1 EMBL,<br>Heidelberg | Titan-Krios IP,<br>Paris | IC-Krios1 EMBL,<br>Heidelberg | Titan-Krios<br>IP, Paris | Glacios IP,<br>Paris |
| Magnification | 165 000 | 105 000 | 165 000 | 105 000 | 150 000 |
| Voltage (kV) | 300 | 300 | 300 | 300 | 200 |
| Electron exposure (e-/Å<sup>2</sup>) | 40 | 40 | 40 | 40 | 50 |
| Defocus range (µm) | -0.6 to -1.6 | -0.6 to -2.2 | -0.6 to -1.6 | -0.6 to -2.2 | -0.8 to -2.4 |
| Pixel size (Å) | 0.731 | 0.86 | 0.731 | 0.86 | 0.96 |
| Symmetry imposed | C5 | C5 | C5 | C5 | C1 |
| Movies | 20151 | 5103 | 11090 | 5757 | 4506 |
| Initial particle images (no.) | 1534810 | 1064223 | 1424083 | 1566807 | 293673 |
| Final particle images (no.) | 104409 | 73690 | 49462 | 114385 | 29644 |
| Map resolution (Å) | 2.3 | 3.4 | 2.7 | 2.8 | 4.4 |
| FSC threshold | 0.143 | 0.143 | 0.143 | 0.143 | 0.143 |
| Map resolution range (Å) | 2.15-8.43 | 3.08-4.22 | 2.5-11.1 | 2.71-7.79 | 4.38-12 |
| **Refinement** |  |  |  |  |  |
| Initial model used (PDB code) |  |  |  |  |  |
| Model resolution (Å) | 2.46 | 3.51 | 2.53 | 2.7 | 3.4 |
| FSC threshold | 0.143 | 0.143 | 0.143 | 0.143 | 0.143 |
| Map sharpening *B* factor (Å<sup>2</sup>) | -100 | -100 | -100 | -100 | -100 |
| Model composition |  |  |  |  |  |
| Non-hydrogen atoms | 13465 | 13615 | 13010 | 13655 | 11145 |
| Protein residues | 1635 | 1635 | 1645 | 1660 | 1393 |
| Ligands | NAG: 20 | NCT:5<br>NAG: 20 | NAG: 20 | NCT: 5<br>NAG: 20 | NAG: 20 |
| *B* factors (Å<sup>2</sup>) |  |  |  |  |  |
| Protein | 72.0 | 78.1 | 38.64 | 84.9 | 29.2 |
| Ligand | 85.4 | 78.5 | 64.14 | 100.0 | 54.4 |
| R.m.s. deviations |  |  |  |  |  |
| Bond lengths (Å) | 0.005 | 0.003 | 0.003 | 0.003 | 0.003 |
| Bond angles (°) | 0.812 | 0.506 | 0.597 | 0.480 | 0.643 |
| Validation |  |  |  |  |  |
| MolProbity score | 2.36 | 2.33 | 2.33 | 1.94 | 2.02 |
| Clashscore | 11.86 | 20.95 | 10.19 | 11.27 | 6.51 |
| Poor rotamers (%) | 4.17 | 3.64 | 3.35 | 1.40 | 0.18 |
| Ramachandran plot |  |  |  |  |  |
| Favored (%) | 95.67 | 91.15 | 93.91 | 96.16 | 85.04 |
| Allowed (%) | 4.33 | 8.85 | 5.97 | 3.84 | 13.36 |
| Disallowed (%) | 0 | 0 | 0.12 | 0 | 1.60 |

**Supplementary Table 2:**
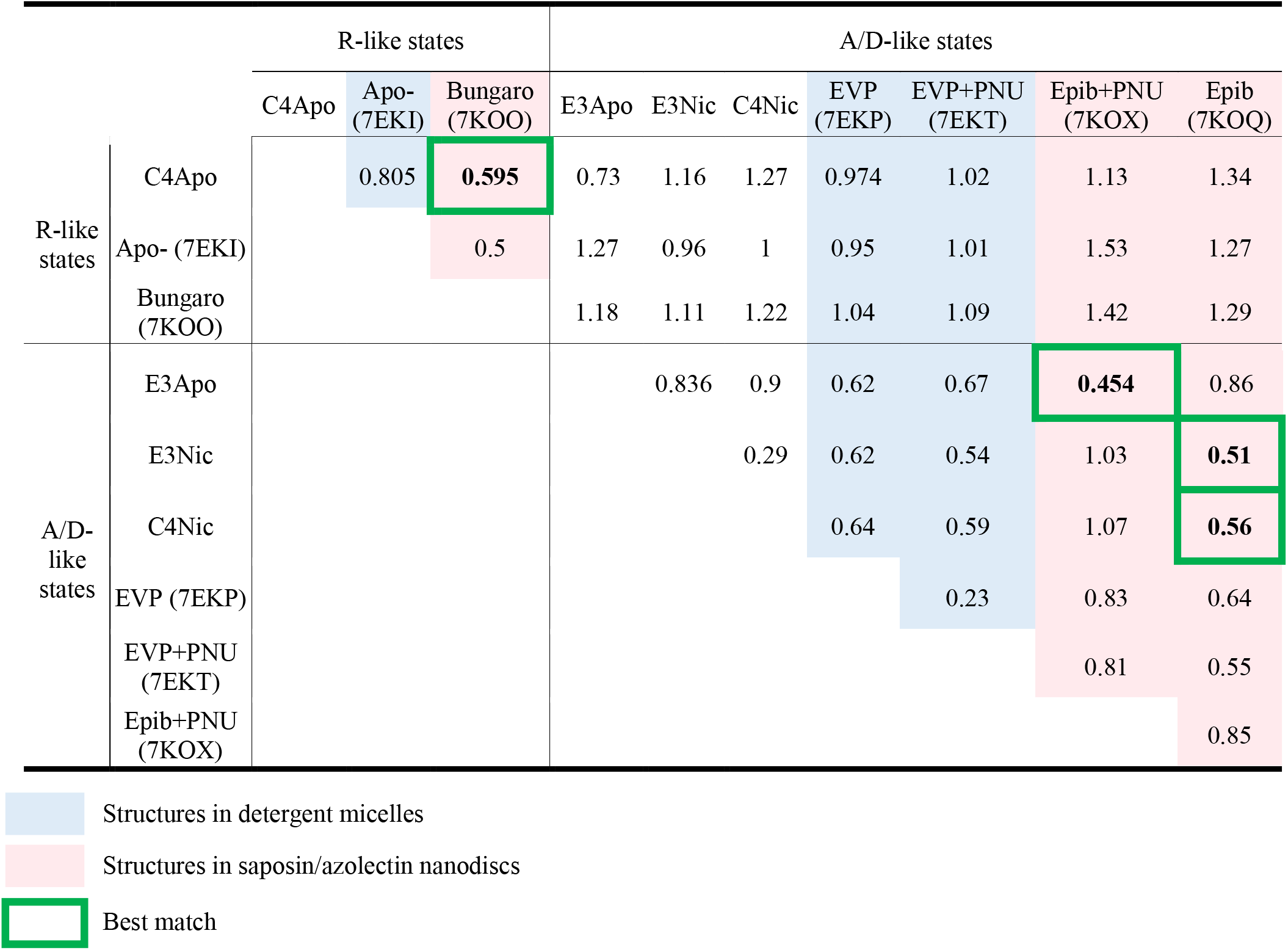
Cα rmsd values calculated on the pentamer of ECD for the indicated structures.

**Supplementary Table 3:**
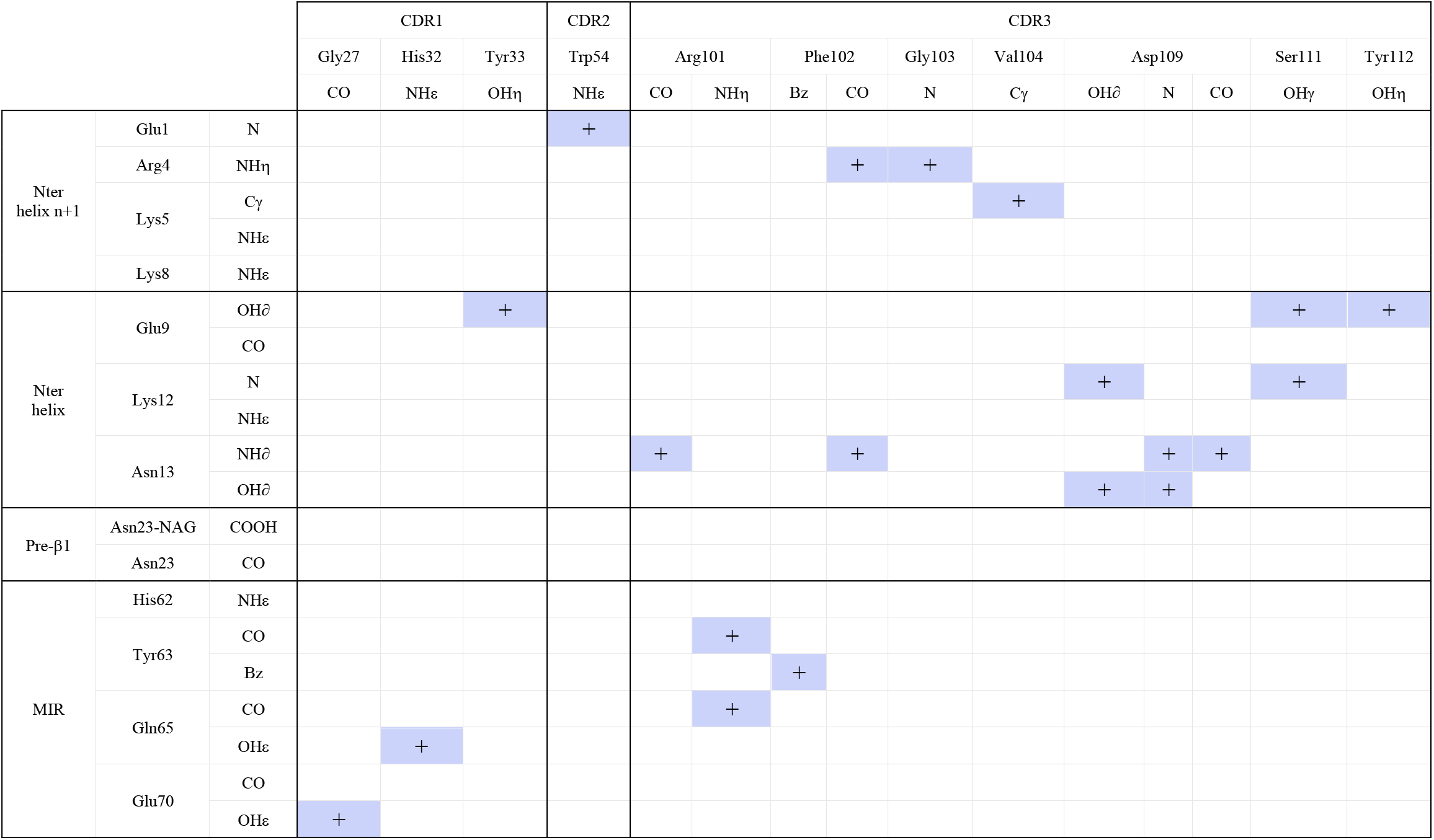
Atomic contacts between C4 and α7. Contacts were selected when the inter-atomic distance is bellow 4 Å, unless specified

**Supplementary Table 4:**
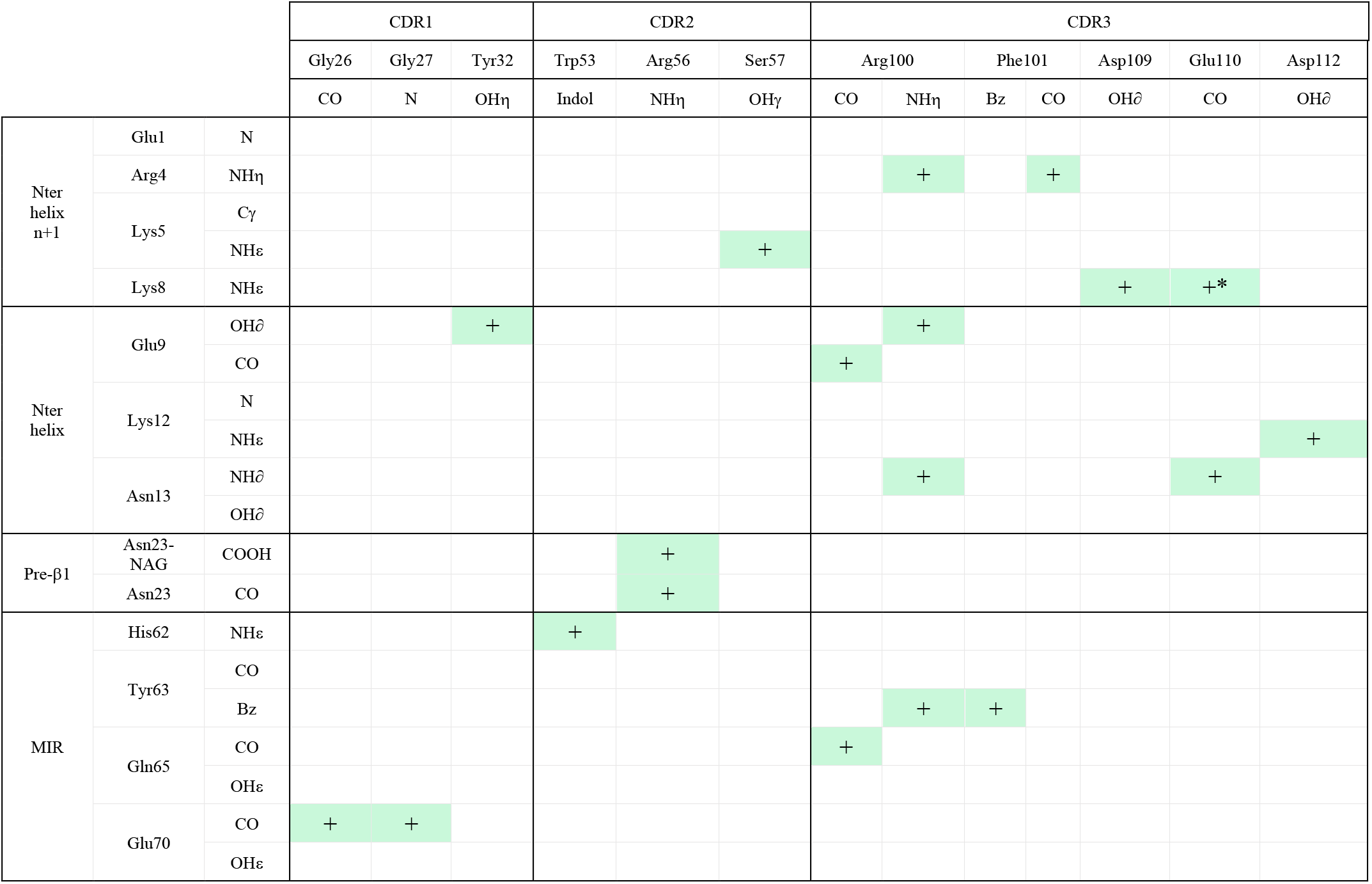
Atomic contacts between E3 and α7. Contacts were selected when the inter-atomic distance is bellow 4 Å, unless specified * 4.4 Å

## References

1. Nemecz, A., Prevost, M. S., Menny, A. & Corringer, P.-J. Emerging Molecular Mechanisms of Signal Transduction in Pentameric Ligand-Gated Ion Channels. Neuron 90, 452–470 (2016).

2. Papke, R. L. & Horenstein, N. A. Therapeutic Targeting of *α* 7 Nicotinic Acetylcholine Receptors. Pharmacol. Rev. 73, 1118–1149 (2021).

3. Hone, A. J. & McIntosh, J. M. Nicotinic acetylcholine receptors in neuropathic and inflammatory pain. FEBS Lett. 592, 1045–1062 (2018).

4. Terry, A. V. α7 nicotinic acetylcholine receptors as therapeutic targets in schizophrenia_ Update on animal and clinical studies and strategies for the future. (2020).

5. Young, G. T., Zwart, R., Walker, A. S., Sher, E. & Millar, N. S. Potentiation of α7 nicotinic acetylcholine receptors via an allosteric transmembrane site. Proc. Natl. Acad. Sci. 105, 14686–14691 (2008).

6. Noviello, C. M. et al. Structure and gating mechanism of the α7 nicotinic acetylcholine receptor. Cell 184, 2121–2134.e13 (2021).

7. Zhao, Y. et al. Structural basis of human α7 nicotinic acetylcholine receptor activation. Cell Res. 1–4 (2021) doi:10.1038/s41422-021-00509-6.

8. Zhuang, Y., Noviello, C. M., Hibbs, R. E., Howard, R. J. & Lindahl, E. Differential interactions of resting, activated, and desensitized states of the α7 nicotinic acetylcholine receptor with lipidic modulators. Proc. Natl. Acad. Sci. 119, e2208081119 (2022).

9. Li, Q., Nemecz, Á., Aymé, G., Prevost, M. S. & Pons, S. Generation of nanobodies acting as silent and positive allosteric modulators of the α7 nicotinic acetylcholine receptor. Preprint at https://www.biorxiv.org/

10. Elegheert, J. et al. Lentiviral transduction of mammalian cells for fast, scalable and high-level production of soluble and membrane proteins. Nat. Protoc. 13, 2991–3017 (2018).

11. Morales-Perez, C. L., Noviello, C. M. & Hibbs, R. E. X-ray structure of the human α4β2 nicotinic receptor. Nature 538, 416–416 (2016).

12. Gu, S. et al. Brain α7 Nicotinic Acetylcholine Receptor Assembly Requires NACHO. Neuron 89, 948–955 (2016).

13. Walsh, R. M. et al. Structural principles of distinct assemblies of the human α4β2 nicotinic receptor. Nature 1–25 (2018) doi:10.1038/s41586-018-0081-7.

14. Gharpure, A. et al. Agonist Selectivity and Ion Permeation in the α3β4 Ganglionic Nicotinic Receptor. Neuron 104, 501–511.e6 (2019).

15. Revah, F. et al. Mutations in the channel domain alter desensitization of a neuronal nicotinic receptor. 353, (1991).

16. Bertrand, D. et al. Unconventional pharmacology of a neuronal nicotinic receptor mutated in the channel domain. Proc. Natl. Acad. Sci. 89, 1261–1265 (1992).

17. Tzartos, S., Langeberg, L., Hochschwender, S. & Lindstrom, J. Demonstration of a main immunogenic region on acetylcholine receptors from human muscle using monoclonal antibodies to human receptor. FEBS Lett. 158, 116–118 (1983).

18. Muyldermans, S. Single domain camel antibodies: current status. Rev. Mol. Biotechnol. 74, 277–302 (2001).

19. Liebschner, D. et al. Macromolecular structure determination using X-rays, neutrons and electrons: recent developments in *Phenix*. Acta Crystallogr. Sect. Struct. Biol. 75, 861–877 (2019).

20. Hassaine, G. et al. X-ray structure of the mouse serotonin 5-HT3 receptor. Nature 1–21 (2014) doi:10.1038/nature13552.

21. Masiulis, S. et al. GABAA receptor signalling mechanisms revealed by structural pharmacology. Nature 1–22 (2018) doi:10.1038/s41586-018-0832-5.

22. Sente, A. et al. Differential assembly diversifies GABAA receptor structures and signalling. Nature 604, 190–194 (2022).

23. Brams, M. et al. Modulation of the Erwinia ligand-gated ion channel (ELIC) and the 5-HT3 receptor via a common vestibule site. eLife 9, 213–19 (2020).

24. Noviello, C. M., Kreye, J., Teng, J., Prüss, H. & Hibbs, R. E. Structural mechanisms of GABAA receptor autoimmune encephalitis. Cell 185, 2469–2477.e13 (2022).

25. Zhu, S. et al. Structure of a human synaptic GABAA receptor. Nature 559, 67–72 (2018).

26. Luo, J. et al. Main Immunogenic Region Structure Promotes Binding of Conformation-Dependent Myasthenia Gravis Autoantibodies, Nicotinic Acetylcholine Receptor Conformation Maturation, and Agonist Sensitivity. J. Neurosci. 29, 13898–13908 (2009).

27. Noridomi, K., Watanabe, G., Hansen, M. N., Han, G. W. & Chen, L. Structural insights into the molecular mechanisms of myasthenia gravis and their therapeutic implications. eLife 6, e23043 (2017).

28. Krause, R. M. et al. Ivermectin: A Positive Allosteric Effector of the ␣7 Neuronal Nicotinic Acetylcholine Receptor.

29. Galzi, J. L., Bertrand, S., Corringer, P. J., Changeux, J. P. & Bertrand, D. Identification of calcium binding sites that regulate potentiation of a neuronal nicotinic acetylcholine receptor. EMBO J. 15, 5824–5832 (1996).

30. Delbart, F. et al. An allosteric binding site of the α7 nicotinic acetylcholine receptor revealed in a humanized acetylcholine-binding protein. J. Biol. Chem. 293, 2534–2545 (2018).

31. Huang, X. et al. Crystal structures of human glycine receptor α3 bound to a novel class of analgesic potentiators. Nat. Struct. Mol. Biol. 24, 108–113 (2016).

32. Eiselé, J. L. et al. Chimaeric nicotinic-serotonergic receptor combines distinct ligand binding and channel specificities. Nature 366, 479–483 (1993).

33. Nury, H. et al. Crystal structure of the extracellular domain of a bacterial ligand-gated ion channel. J. Mol. Biol. 395, 1114–1127 (2010).

34. Moraga-Cid, G. et al. Allosteric and hyperekplexic mutant phenotypes investigated on an α 1glycine receptor transmembrane structure. Proc. Natl. Acad. Sci. U. S. A. 201417864 (2015) doi:10.1073/pnas.1417864112.

35. Hu, H., Howard, R. J., Bastolla, U., Lindahl, E. & Delarue, M. Structural basis for allosteric transitions of a multidomain pentameric ligand-gated ion channel. Proc. Natl. Acad. Sci. 117, 13437–13446 (2020).

36. Lykhmus, O. et al. α7 Nicotinic Acetylcholine Receptor-Specific Antibody Induces Inflammation and Amyloid β42 Accumulation in the Mouse Brain to Impair Memory. PLOS ONE (2015).

